# Orphan GPCRs as Targets for Human Brain Modulation: A Multi-omic Atlas of Cell-Type Specific Expression

**DOI:** 10.64898/2026.03.23.713764

**Authors:** Alan Umfress, Patrick Wertimer, Christina Pressl, Matthew Baffuto, Kert Mätlik, Franchesca Fernandez, Maria Esterlita Siantoputri, Ines Ibañez-Tallon, Nathaniel Heintz

## Abstract

G Protein coupled receptors (GPCRs) are the largest class of clinically validated drug targets with nearly 35% of all approved therapeutic agents acting on these receptors. To further explore the potential of this class of receptors for the development of circuit-specific and mechanism-based therapeutic strategies for neurological disorders, we focused on GPCRs with no known endogenous ligand, orphan GPCRs (oGPCRs), because knowledge of their functions in the human brain remains rudimentary. Here, we utilized fluorescence activated nuclear sorting and sequencing (FANSseq) to generate deep molecular profiles of cell type specific nuclei isolated from post-mortem brains to generate an atlas of oGPCR expression across multiple regions of the human brain. We identified 22 oGPCRs that displayed selective cell-type enrichment both in RNA transcript expression and chromatin accessibility. We further validated each of these targets for cell-type specific expression in human brains and developed an open-source web atlas of all oGPCR expression in the human brain to serve as a neuro-resource for the broader scientific community. These studies reveal novel cell-type specific expression patterns of several oGPCRs, suggest potential endogenous roles for these receptors, and identify validated candidates for cell-type specific neuromodulation of the human brain.

**One Sentence Summary:** This study presents an atlas of orphan GPCR expression across the human brain for translational targeting.

## INTRODUCTION

G protein-coupled receptors (GPCRs) mediate a diverse range of physiological processes within the central nervous system through their ability to transduce extracellular signals into intracellular responses (*1*). GPCRs play central roles in modulating neurotransmission, neuroplasticity, neuroinflammation, and neurodevelopment (*1, 2*). Dysregulation in GPCR signaling has been implicated in a spectrum of neurological and psychiatric disorders, including schizophrenia, depression, Alzheimer’s disease (AD), Parkinson’s disease (PD), and epilepsy (*2–4*). Due to these expansive roles in regulating cellular function, GPCRs are pivotal targets in pharmacology, with more than one-third of all FDA-approved drugs acting on GPCRs (*5*). Decades of research have uncovered vast amounts of information across diverse sets of GPCRs including, their cell-type expression patterns, high-resolution electron microscopy structures, and novel pharmacological agents to target these receptors in diseases (*2*). This translational targeting of GPCRs is especially pertinent in diseases of the central nervous system (CNS) where select agonism, antagonism, and signaling biases of GPCRs allow for the selective tuning of neural excitability and circuit dynamics.

Despite the remarkable success in GPCR research, a substantial group of receptors remain classified as orphan, lacking a known endogenous ligand (*6*). Most class A oGPCRs are expressed in the brains of mammals, yet for many receptors no CNS function has been described (*6*). By defining which cells and circuits oGPCRs modulate within the brain, we can begin to narrow down their endogenous functions and ligands, illuminating their therapeutic potentials. For a small subset of oGPCRs with well-defined cell-type expression, therapeutic agents targeting these receptors have translated into clinical trials (*7, 8*). For example, defining the cell-type specific expression of the orphan receptor GPR6 in indirect medium spiny projection neurons (iMSNs) of the striatum led to the development of selective inverse agonist targeting of GPR6 in PD to modulate iMSN projections that mediate symptomatic dyskinesia (*8–11*). The trials for the inverse agonist modulating GPR6 have advanced into Phase III. Furthermore, elucidation of GPR139 expression within the medial habenula, lateral habenula, and the striatum has yielded trials assessing this receptor’s therapeutic potential in major depressive disorder and schizophrenia (*12–15*). Additionally, non-CNS targeting of oGPCRs such as GPR119 in enteroendocrine and pancreatic islet cells is being tested as a positive regulator of glucagon-like peptide 1 (GLP-1) for the treatment of diabetes (*16*). Hence, these receptors represent a broad and untapped target class for the development of new translational treatments.

To define the cell type expression of oGPCRs across the human brain for translational study, we have used fluorescence activated nuclei sorting and sequencing (FANSseq) data to generate an atlas of oGPCR expression across multiple regions of the human brain including the neocortex, the striatum, the cerebellum, and the hippocampus. We have taken deep-molecular profiles across these regions to define gene expression patterns of all class A oGPCRs across 27 distinct cell-types, enabling the identification of receptors with cell-type specific expression. To this end, we created a web-application to serve as a neuro-resource for the broader scientific community (https://cpressl.shinyapps.io/GPCRxplorer/), enabling exploration of class A oGPCRs across human brain cell-types. We then validated and filtered cell-type specific oGPCR expression by assessing the chromatin accessibility of each receptor within the cell-types using ATAC-seq data. Ultimately, this approach identified 22 Class A oGPCRs with cell-type specific or enriched expression across brain regions and cell-types of the human brain. This analysis revealed both conserved expression patterns of oGPCRs between humans and mice, as well as distinct human specific expression within the CNS. Finally, we validated the cell-type expression patterns of each selected oGPCR using cell-type specific markers and *in situ* hybridization in post-mortem human brains. Defining the cell- and circuit-specific expression patterns of these oGPCRs is the first step towards understanding their physiological roles and developing strategies that target individual oGPCRs for the treatment of neurological diseases.

## RESULTS

### Cell-type atlas of class A oGPCRs in the human brain

To generate an atlas of oGPCR expression we utilized available molecular profiles and generated novel FANSseq profiles of cell types within the human cortex, hippocampus, striatum, and cerebellum (Fig. 1A) (*17–19*). FANSseq yields deep molecular profiles of approximately 12,000-15,000 genes per cell type, enabling detection of GPCR targets that are often expressed at very low levels in brain cell types. FANSseq of human hippocampal nuclei was done by separating nuclei of neuronal and non-neuronal cell types, with neuronal cells segmented into three populations of excitatory neurons (ENs) or one population of Interneurons (INs). ENs were regionally divided into granule neurons of the dentate gyrus (hGN), *Cornu ammonis* 1 (CA1), and *Cornu ammonis* 2/3 (CA2/3), while INs were collected in one population of Adenosine Deaminase RNA Specific B2 ADARB2^+^ cells (Fig. 1B). Non-neuronal cell-types from the hippocampus were separated into four populations including astrocytes, microglia, oligodendrocytes, and oligodendrocyte precursor cells (OPCs). Cells of the human striatum were also divided into neuronal or non-neuronal cell types. As previously demonstrated(*18*), striatal cell types analyzed included striatonigral direct projecting medium spiny neurons (dMSNs), striatopallidal indirect projecting medium spiny neurons (iMSNs), interneurons expressing Tachykinin Precursor 3 (TAC3+), Somatostatin (SST), Choline Acetyltransferase (CHAT+) or Parvalbumin (PVALB+), and the four glial astrocytes, microglia, oligodendrocytes, and OPCs. Similarly, cell populations of human cerebellum samples analyzed were cerebellar granule neurons (cGN), basket cells (BC), and Purkinje Cells (PC), as well as astrocytes, microglia, oligodendrocytes, and OPCs. Neocortical cell types studied here include pyramidal cells from cortical layers 2/3, 4, 5a, 6a, and 6b, Von Economo neurons (VENs), Betz cells, four populations of interneurons including those expressing PVALB+, Vasoactive Intestinal Peptide (VIP+), Reelin+, and LAMP5+, and the non-neuronal populations of astrocytes, microglia, oligodendrocytes, and OPCs (*20*). Taken together this FANS-seq data is comprised of deep molecular profiles of 27 principal cell types, as well as region specific non-neuronal glial populations from four distinct brain regions.

**Figure 1.**
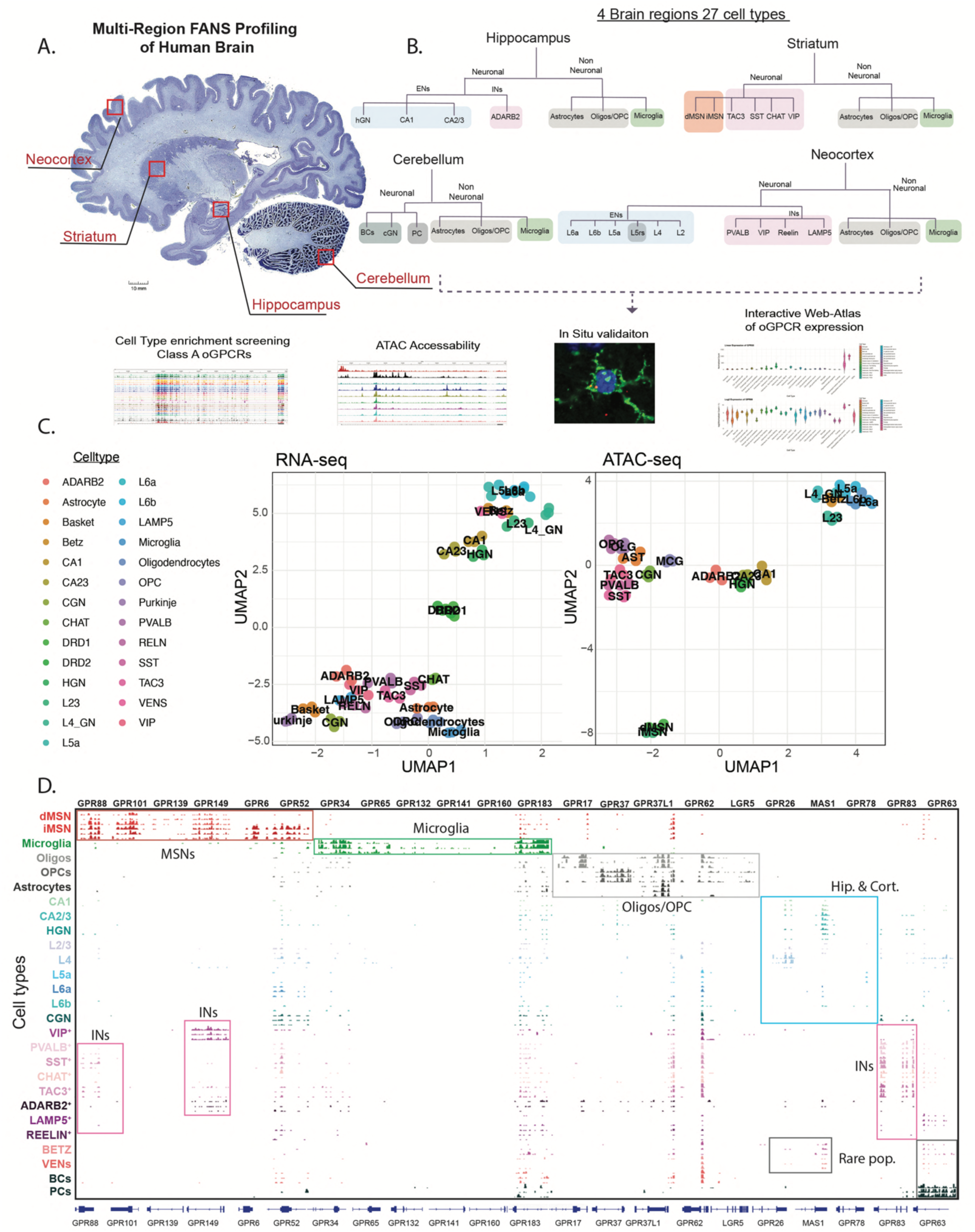
FANS-seq profiling of oGPCR cell type or brain region enriched oGPCR expression in the human brain. A. Human brain schematic depicting brain regions and studies used for cell profiling (image adapted from: brainmuseum.org). B. Flow chart representing the 4 brain regions and cell types assessed for oGPCR expression (top) and follow-up validation to confirm cell-type expression within the human brain (bottom). C. UMAPs displaying cell type clustering for all 27 cell-types assessed by FANS-RNA-seq (left) and FANS-ATAC-seq (right) across 21 cell-types. D. IGV plot of FANS-seq data displaying 22 identified cell-type enriched oGPCRs within the human brain including subgroup specific oGPCRs selective for MSNs (red box), microglia (green box), oligodendrocytes and oligodendrocyte precursor cells (grey box), Excitatory neurons of the hippocampus and cortex (blue box), Interneurons (purple box), and rare cell populations including Von Economo Neurons, Betz cells, and Purkinje cells (black box) each cell type has 3 representative tracks.

Utilizing the FANSseq datasets, we screened all Class A oGPCRs for cell-type specific mRNA expression across the human brain, assessed each gene for chromatin accessibility, validated the cell-type specific expression via RNAscope, and generated a searchable atlas of oGPCR expression in the human brain (https://cpressl.shinyapps.io/GPCRxplorer/) (Fig. 1B). Uniform manifold approximation projection analysis (UMAP) of FANSseq data from each brain region and cell-type resolved into in distinct clusters that included neocortical excitatory cell types, hippocampal excitatory neurons, direct and indirect projecting MSNs, and distinct classes of interneurons, cerebellar neurons, and non-neuronal cells (Fig. 1C). Likewise, cells for which ATAC-seq was also performed showed similar separation among the four brain regions and cell types (Fig. 1C). Screening of all Class A oGPCRs for cell type specific enriched expression revealed 22 orphan receptors with unique expression profiles clustered into main cell-type groups including oGPCRs expressed in MSNs, microglia, Oligodendrocytes and OPCs, ENs of the hippocampus and neocortex, and receptors expressed in rare cell populations such as VENS, Betz, and Purkinje cells (Fig. 1D). We identified six receptors with selective expression in MSNs, with four oGPCRs showing expression in both dMSNs and iMSNs (*GPR101, GPR88, GPR139, and GPR149*), and two receptors showing expression in only the iMSNs (GPR6 and GPR52) (Fig. 1D). Similarly, six oGPCRs were identified with selective expression in human microglia (*GPR32, GPR65, GPR132, GPR141, GPR160,* and *GPR183)*. We also identified five oGPCRs with cell-type enrichment in either oligodendrocytes or OPCs including *GPR37, GPR17, GPR37L1, GPR62,* and *LGR5.* Interestingly, *GPR17* showed much higher expression in OPCs compared to its expression in mature oligodendrocytes. Additionally, GPR37L1 shared expression in astrocytes in addition to oligodendrocytes and OPCs. While no oGPCR showed enrichment in a single hippocampal or cortical neuronal population, we identified *GPR26, MAS1,* and *GPR78* as oGPCRs enriched in ENs of the hippocampus and cortex, with *GPR78* displaying the most selectivity for L5a pyramidal neurons. Additionally, expression of *GPR26* and *MAS1* was detected in rare cortical populations such as VENS and Betz cells. *GPR63* expression was detected in several layers of the neocortex, as well as, in VENs and Betz cells. However, *GPR63* expression was significantly higher in Purkinje cells than any other cell-type examined. *GPR83* was the only oGPCR we observed with expression mainly restricted to striatal interneuron populations such as VIP+, SST+, CHAT+, and TAC3+ cells. However, some receptors with expression in inhibitory MSNs also displayed enrichment in inhibitory interneuron populations of the human striatum including *GPR149*, which was largely detected in VIP+ interneurons and *GPR88* which showed broad expression in interneurons populations. Altogether, these data reveal 22 oGPCRs as potential drug targets in the human brain for selective cell-type neuromodulation.

While the identified orphan receptors may be used for selective cell-type targeting, many other oGPCRs screened showed expression in the human brain, while some showed no detectable expression (Fig. S1A). For example, *GPR176*, which is implicated in interneuron function, cancer, and circadian rhythm modulation (*21–23*) shows broad expression in all cortical excitatory neurons and interneuron populations (Fig. S1B). In contrast, some families of oGPCRs such as the trace-amine receptors like TAAR5 showed no expression across these human CNS cell-types (Fig. S1A,C). This could suggest no CNS expression of these receptors, or localization to another brain area not assessed here. For example, immunohistochemistry of the mouse and human habenula shows an evolutionarily conserved cell-type selective expression of the orphan receptor *GPR151* in the habenula and its projections to the interpeduncular nucleus (*24*) (Fig. S1D). Collectively, these data show many oGPCRs are expressed in a cell-type-specific manner in the human brain and may play crucial roles in CNS function.

Vast amounts of molecular profiling of cell-types across species has occurred in recent years yielding the generation of several databases of cell-type gene expression patterns (*25–27*). Here, we utilized FANS-seq data (*17–19*) to define oGPCR expression across the human brain due to its ability to detect lowly expressed targets (*28*) and further validated these finding by inspection of ATAC-accessibility within each cell-type. For comparative analysis, we also assessed publicly available databases of human cortex using SMART-seq and 10x single-cell sequencing from the Allen Brain (Fig. S2). We observed little to no expression of these oGPCR targets in the available SMART-seq data (Fig. S2A). However, 10x single-cell was able to detect cell-type expression of a few of these oGPCR targets including GPR149 expression in interneuron populations, GPR63 expression throughout cortical layers, and oligodendrocyte specific expression of GPR37 (Fig S2B), while other oGPCRs remained undetected.

The search for endogenous ligands and the underlying roles for many of these receptors within these CNS cell-types is ongoing. Structural studies of many ligand-elusive oGPCRs suggest GPCR activity may be ligand-independent via GPCR constitutive activation (*29, 30*). Interestingly, screens of constitutive GPCR and oGPCR activity reveal many of the highest constitutively active GPCRs to be orphans in a β-arrestin recruitment assay (*31*) (Fig. S3A). Likewise, many of the 22 receptors identified as cell-type enriched in the CNS show high constitutive activity (Fig. S3B). Examples include GPR101, GPR37, and GPR52 all of which have been shown to have conserved second extracellular loop (ECL2) domains that mediate their high basal activity and block the traditional orthosteric ligand binding pocket. Whether these receptors are solely genetically regulated or also respond to stimulation of their cell-type circuits, defining each receptor’s regional and cell-type localization will aid in discovering their ligands, potential functions, and therapeutic implications.

### Identifying oGPCR targets for striatal neuromodulation

Modulation of the striatonigral and striatopallidal pathways has been implicated in numerous neurological conditions, including addiction, stress, anxiety, obsessive-compulsive disorder, and PD (*32–35*). Thus, novel pharmacological approaches modulating the activity of these circuits via GPCR targeting remains pivotal for therapeutic development. To expand the druggable targets comprising these brain circuities, we utilized FANS-ATAC-seq datasets of chromatin accessibility to validate the RNA expression patterns of the six MSN specific oGPCRs identified. In agreement with FANS-seq based transcript data, we observed that six oGPCRs, including *GPR88, GPR101, GPR139, GPR149, GPR6,* and *GPR52* contained chromatin regions highly accessible in MSNs, indicative of stronger recruitment of potential enhancers in these neurons (Fig. 2A). Additionally, *GPR6* and *GPR52* displayed cell-type specific accessible peaks within iMSNs while *GPR88, GPR101, GPR139* and *GPR149* exhibited accessible peaks in both dMSNs and iMSNs (Fig. 2A). Next, to examine oGPCR expression in the human brain, we performed cell-type-specific peak-to-gene annotation through integration of the FANS-seq RNA expression data with ATAC-seq consensus peaks using a correlation-based linkage approach. This allowed us to visualize a correlation between the expression of each oGPCR and the accessibility of each oGPCR in all 27 cell types. As expected, the six MSN specific oGPCRs showed a high correlation between gene accessibility in MSNs and transcript level (Fig. 2B). For quantitative visualization of each receptor’s RNA expression, sequencing depth-normalized number of FANS-seq reads mapped to each oGPCR gene were plotted. For example, *GPR139* shows significantly higher expression in dMSNs and iMSNs relative to all other cell types. Likewise, *GPR6* expression is restricted to iMSNs (Fig. 2C). To further validate the FANS-seq datasets, we confirmed the expression of each striatal oGPCR in the striatum of human brain donors using RNAscope, showing that each receptor’s expression co-localized with the well-established MSN marker DARPP-32 (*36*). We observed cell-type specific staining of *GPR101, GPR88, GPR139,* and *GPR149* within all DARPP32 positive MSNs (Fig. 2D). Alternatively, we observed high expression of GPR6 in some DARPP32 positive MSNs and no expression in others, in agreement with FANS-seq data showing GPR6 expression is restricted to iMSN where it could modulate the activity of the indirect pathway. Collectively, this dataset provides a comprehensive characterization of striatal oGPCR expression and highlights how these targets may be useful for modulating dopaminergic signaling within the direct vs indirect pathways in neurological disease.

**Figure 2.**
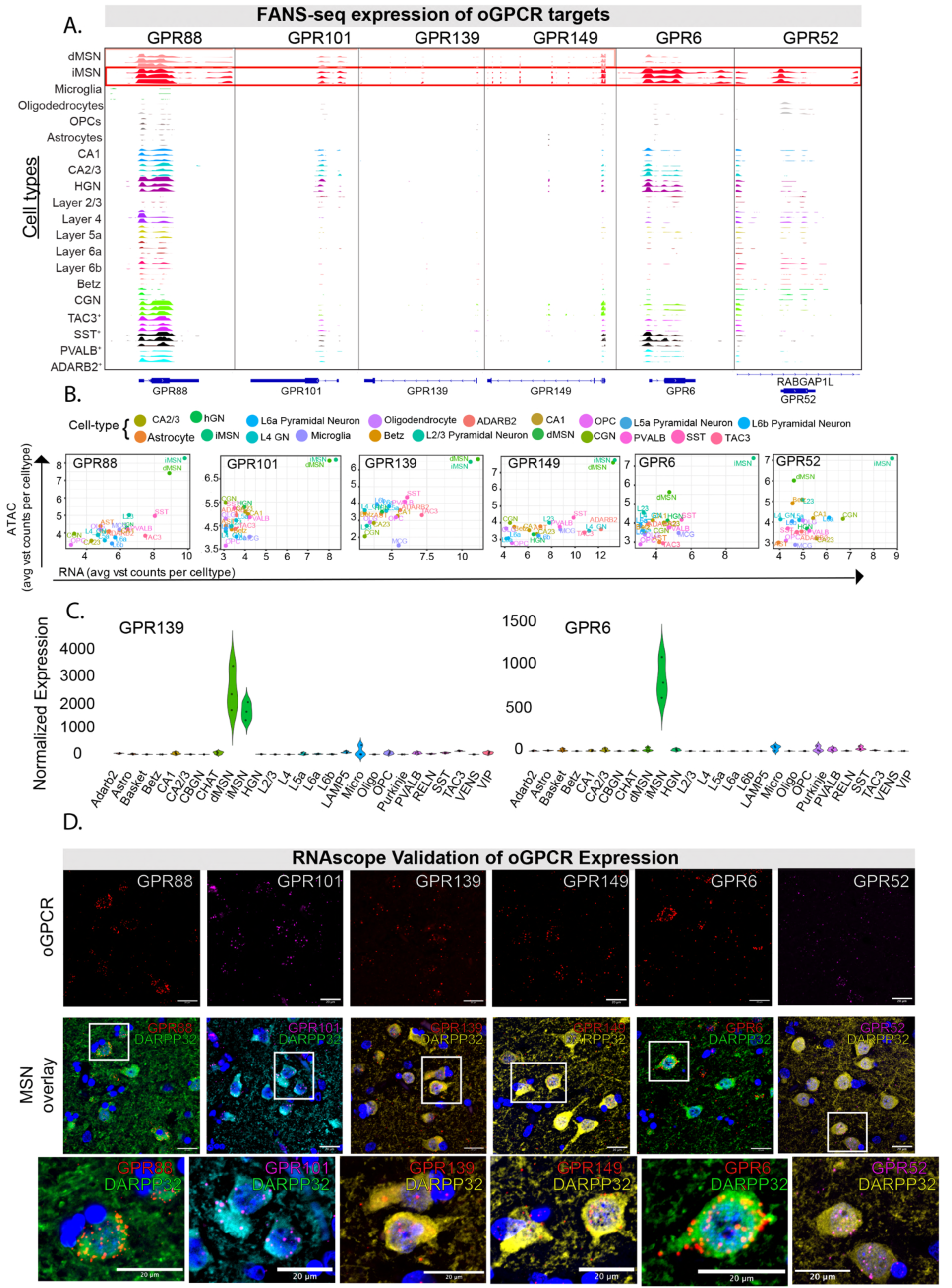
Accessibility and expression of selected oGPCR genes in striatal projection neurons. A. IGV plot of ATAC-seq reads’ distribution over 6 oGPCR genes with enriched expression in striatal MSNs, showing chromatin accessibility across human brain cell types (3 tracks per cell type). B. Correlation analysis between ATAC-accessibility and RNA expression of each oGPCR displaying enrichment in MSNs. C. Violin plots from the developed Web-atlas of oGPCR expression depicting RNA expression of *GPR139* and *GPR6* showing selective enrichment in dMSNs and iMSNs of the human brain. D. RNAscope-based detection of oGPCR transcripts in sections of human caudate overlayed with the signal from immunostaining of MSN-specific marker DARPP32.

In addition to the oGPCR enrichment we observed in the human striatum, FANS profiling allowed the detection of lowly expressed receptors in other cell types, including several interneuron populations of the striatum. While receptors such as *GPR149* and *GPR88* showed the highest enrichment in MSNs, we also observed weak expression and accessibility of these receptors in interneuron subtypes (Fig. 2B). For example, *GPR149*, also showed expression in hippocampal interneuron populations. To investigate this, we co-labeled human hippocampal sections with *GPR149* with GABAergic marker GAD2 and hippocampal interneuron marker ADARB2 and confirmed *GPR149* expression in interneuron populations of the hippocampus (Fig. S4A, B). Likewise, FANS-profiling suggested weak *GPR88* expression in interneuron populations of the striatum (Fig. 2B). To confirm this, we performed RNAscope of human caudate to examine *GPR88* expression within SST+ populations of interneurons. Consistent with the FANS-profiling, *GPR88* expression was detected within SST+ interneurons (Fig. S4C). These data highlight how deep profiling using FANS-seq allows us to determine both cell-types with enriched receptor expression and cell-types with lowly expressed GPCR targets.

### Identifying microglia-selective oGPCRs for CNS immune modulation

Microglia serve as the principal immune cells of the CNS, responding to both anti-and pro-inflammatory signaling molecules from the surrounding microenvironment (*37, 38*). Well established GPCR signaling governs microglial state changes, phagocytotic ability, inflammatory capacity, and the progression of neurodegenerative disease (*39, 40*). Thus, microglia and their underlying intracellular signaling states have been implicated in aging and nearly every neurogenerative disease (*37, 41*). However, less is known about oGPCR regulation of microglia with only a few targets assessed for preclinical therapeutic intervention in the CNS (*42*). To identify potential oGPCRs expressed in microglia as potential modulators of neuroinflammation and disease we examined the FANS-ATAC-seq dataset to assess whether oGPCR genes expressed specifically in microglia also exhibit chromatin accessibility specific to this cell type. All six receptors including *GPR183, GPR34, GPR65, GPR132, GPR160,* and *GPR141* showed selective chromatin accessibility in microglia (Fig. 3A). Furthermore, correlations of ATAC accessibility and transcript expression displayed a robust enrichment of all six receptors in microglia (Fig. 3B). This analysis also unveiled a several folds lower expression pattern of GPR141 in oligodendrocytes (Fig. 3B). Gene counts from FANS-seq displayed a strong enrichment of these receptors within microglia including *GPR34* and *GPR132* (Fig. 3C). Finally, RNAscope validation in human BA9 neocortex showed cell-type specific expression of all six oGPCRs within ionized calcium-binding adaptor molecule 1 (IBA1+) microglia marker (*43*) (Fig. 3D).

**Figure 3.**
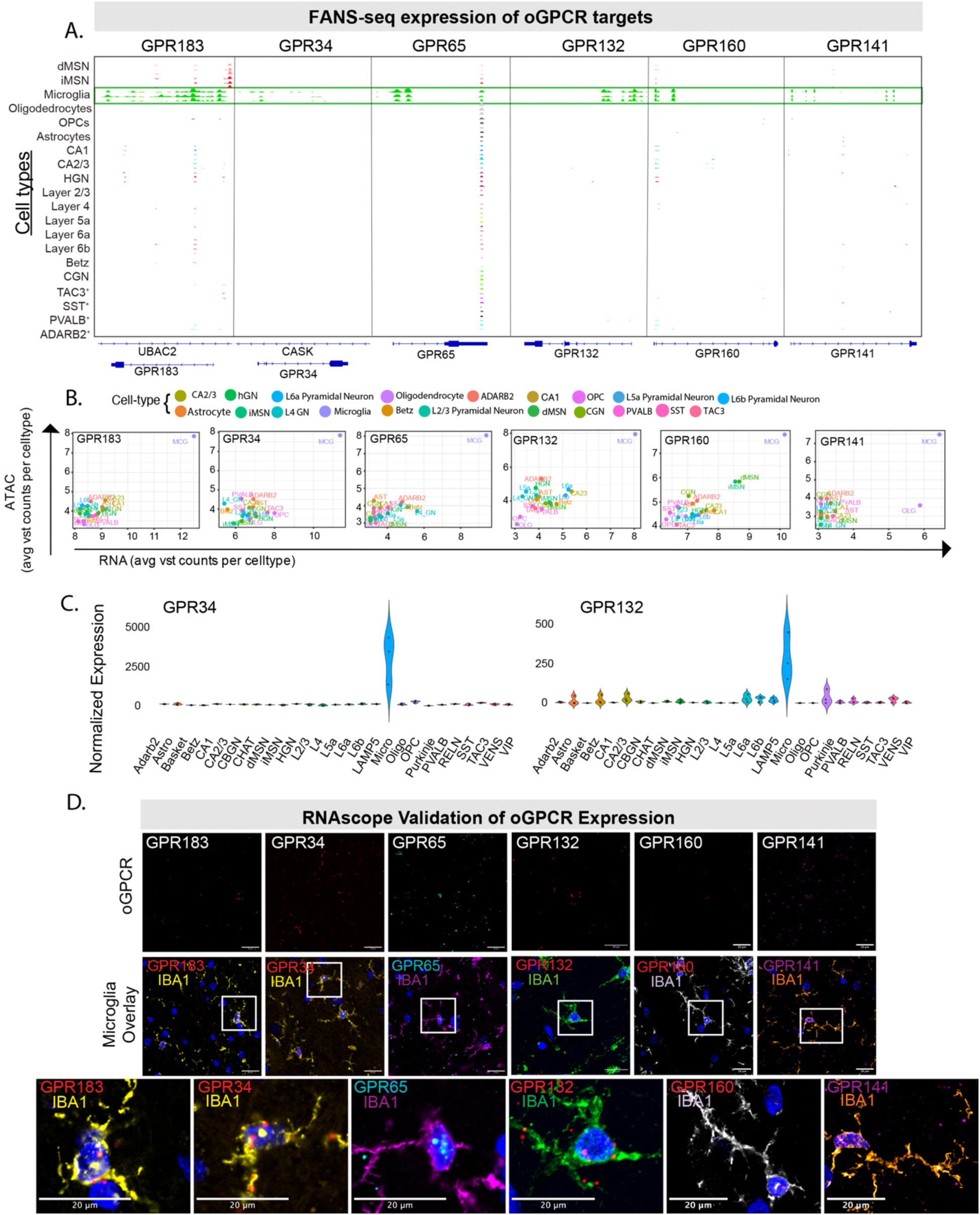
Accessibility and expression of selected oGPCR genes in microglia. A. Distribution of cell type-specific ATAC-seq reads over oGPCR genes with enriched expression in microglia. B. Correlation analysis between chromatin accessibility and RNA expression of each oGPCR displaying enrichment in microglia. C. Violin plots from developed resource atlas depicting expression of *GPR34* and *GPR132* in microglia of the human brain. D. RNAscope-based detection of oGPCR transcripts on sections of human hippocampus and neocortical region BA4, overlayed with the signal from immunostaining of microglia-specific marker IBA1.

### Oligodendrocyte-enriched oGPCR targets

To define oGPCRs regulating oligodendrocyte and OPC specific cell-signaling, we assessed the chromatin accessibility over each of the five cell-enriched oGPCR genes (Fig. 4A). All five oligo/OPC receptors displayed accessible ATAC peaks within gene bodies and at their promoters. Additionally, *GPR37L1* also showed near equivalent accessibility and expression in astrocytes relative to oligos/OPCs (Fig. 4 A,B). This is expected as dual cell-type roles for *GPR37L1* are evolutionarily conserved and have been implicated in astrocytic regulation of pain and oligodendrocyte maturation (*44, 45*). However, RNA-ATAC correlation matrices revealed very selective expression of *GPR62, GPR37,* and *GPR17* to oligo and OPC cell-types (Fig. 4 B,C). To validate the cell-type specific expression patterns of these receptors, we co-localized mRNA probes for each receptor with the cell-type marker OLIG2, which labels both mature oligodendrocytes and OPCs (*46*) (Fig. 4 D). In all cases OLIG2-positive cells expressed the corresponding oGPCR within the human cortex. Remarkably, we also observed nearby GFAP+ astrocytic labeling of *GPR37L1* alongside an adjacent OLIG2+ oligodendrocyte, demonstrating the expression of this receptor in both cell-types of the human brain (Fig. 4 B,D). Here, we define oligodendrocyte and OPC enriched oGPCRs in the human brain as potential targets for demyelinating diseases and highlight potential oGPCRs for modulation of immature or mature oligodendrocyte lineages.

**Figure 4.**
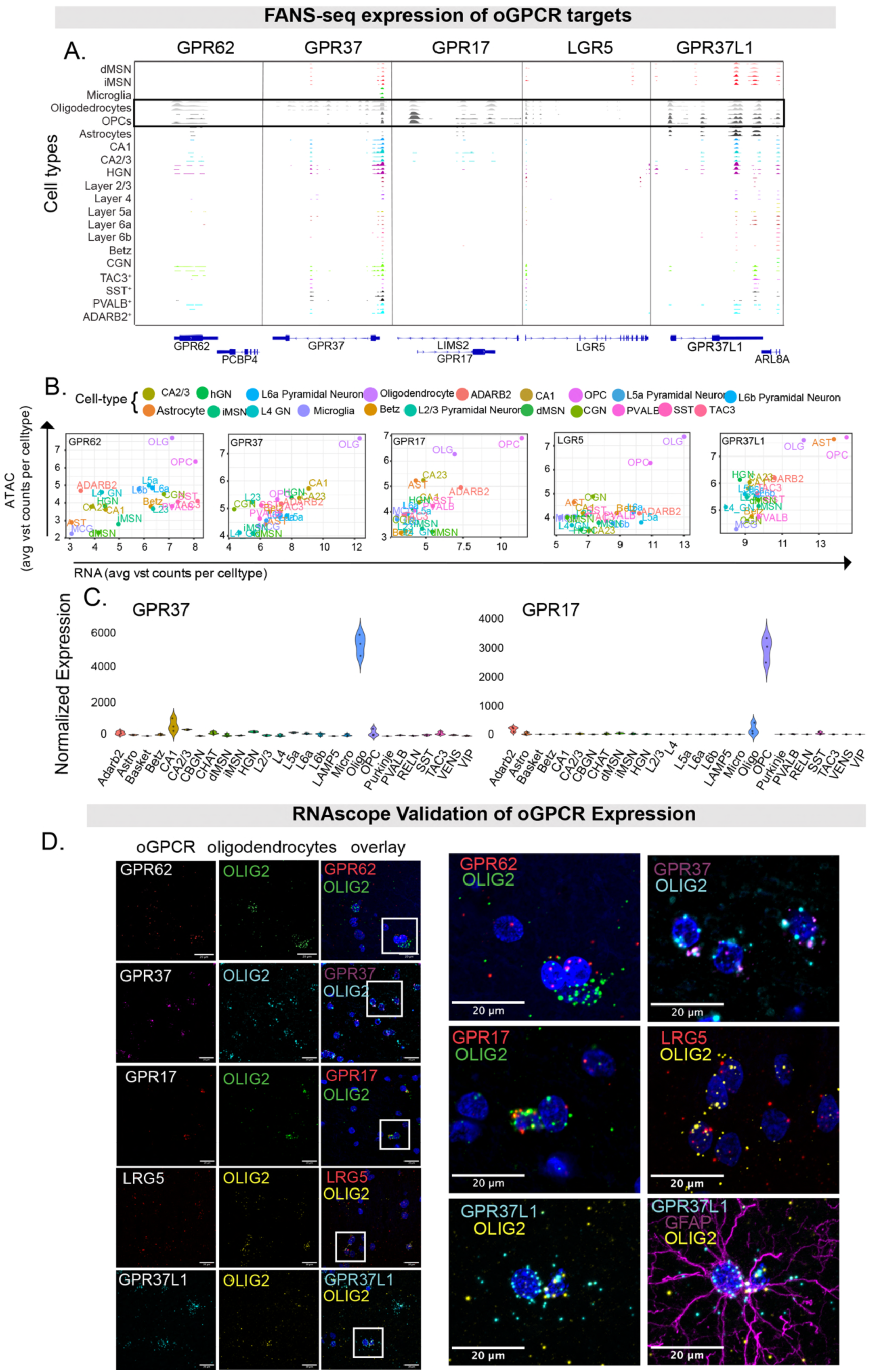
Accessibility and expression of selected oGPCR genes in oligodendrocytes, OPCs, and astrocytes. A. Distribution of cell type-specific ATAC-seq reads over 5 oGPCR genes with enriched expression in oligodendrocyte lineage. B. Correlation analysis between ATAC-accessibility and RNA expression of each oGPCR displaying enrichment in oligos/OPCs. C. Web-atlas RNA expression of *GPR37* and *GPR17* showing selective enrichment in dMSNs and iMSNs of the human brain. D. RNAscope-based detection of oGPCR transcripts on sections of human hippocampus and neocortical region BA4, overlayed with the signal from the transcripts of oligodendrocyte lineage-specific marker gene *OLIG2*.

### oGPCR enrichment in neocortical and hippocampal excitatory neurons

The principal excitatory neurons (ENs) of the hippocampus and neocortex play a crucial role in cognitive function. These neurons mediate perception, cognition, memory, and behavior across species. Altered activity or impairments in these circuits underlie neurological diseases spanning neurodevelopment to neurodegeneration (*47, 48*). Importantly, these cells and their district projections display cell-type specific vulnerability in numerous diseases, such as the early degeneration of CA1 hippocampal pyramidal neurons in Alzheimer’s disease and the vulnerability of deep layer neocortical neurons in Huntington’s disease (*19, 49*). Thus, molecular modulators of these cells may represent disease modifying signaling pathways to perturb disease progression. To identify oGPCRs with cell-type specific expression in ENs of the cortex and hippocampus we cross-referenced our ATAC-seq data with the FANS-RNA-seq profiled cells to search for selective expression of oGPCR targets. While we observed no singular cell-type specific oGPCRs to the 3 EN populations of hippocampal neurons or across cortical layers, high chromatin accessibility confirmed broadly EN-enriched expression of 3 oGPCR genes including *MAS1, GPR26,* and *GPR78* (Fig. 5A). Correlating the expression to accessibility of each receptor we observed pyramidal cell enrichment of all 3 oGPCRs within the cortex and hippocampus, with weaker expression of *MAS1* and *GPR26* also observed within ADARB2+ interneurons of the hippocampus (Fig. 5B). To confirm the expression of oGPCR targets in human brain tissue, we performed RNAscope for each oGPCR assessing co-localization with EN markers. We observed expression of *MAS1* co-localized with EN marker Solute Carrier Family 17 Member 6 (SLC17a7, VGLU1) within the CA1 pyramidal cell layer of the hippocampus. Additionally, *GPR26* co-localized with 17a7 within ENs of the cortex (BA9) (Fig. 5C). *GPR78* showed the most cell-specific enrichment in deep-layer cortical neurons particularly in L5a (Fig. 5 A,B). To confirm this layer-enrichment, we colocalized *GPR78* expression with Purkinje cell protein 4 (*PCP4*), a marker of cerebellar Purkinje cells that also shows selective expression in deep layers of the human cortex (*50*) (Fig. 5C). Together, these results highlight 3 potential neuromodulators of the main ENs within the human brain.

**Figure 5.**
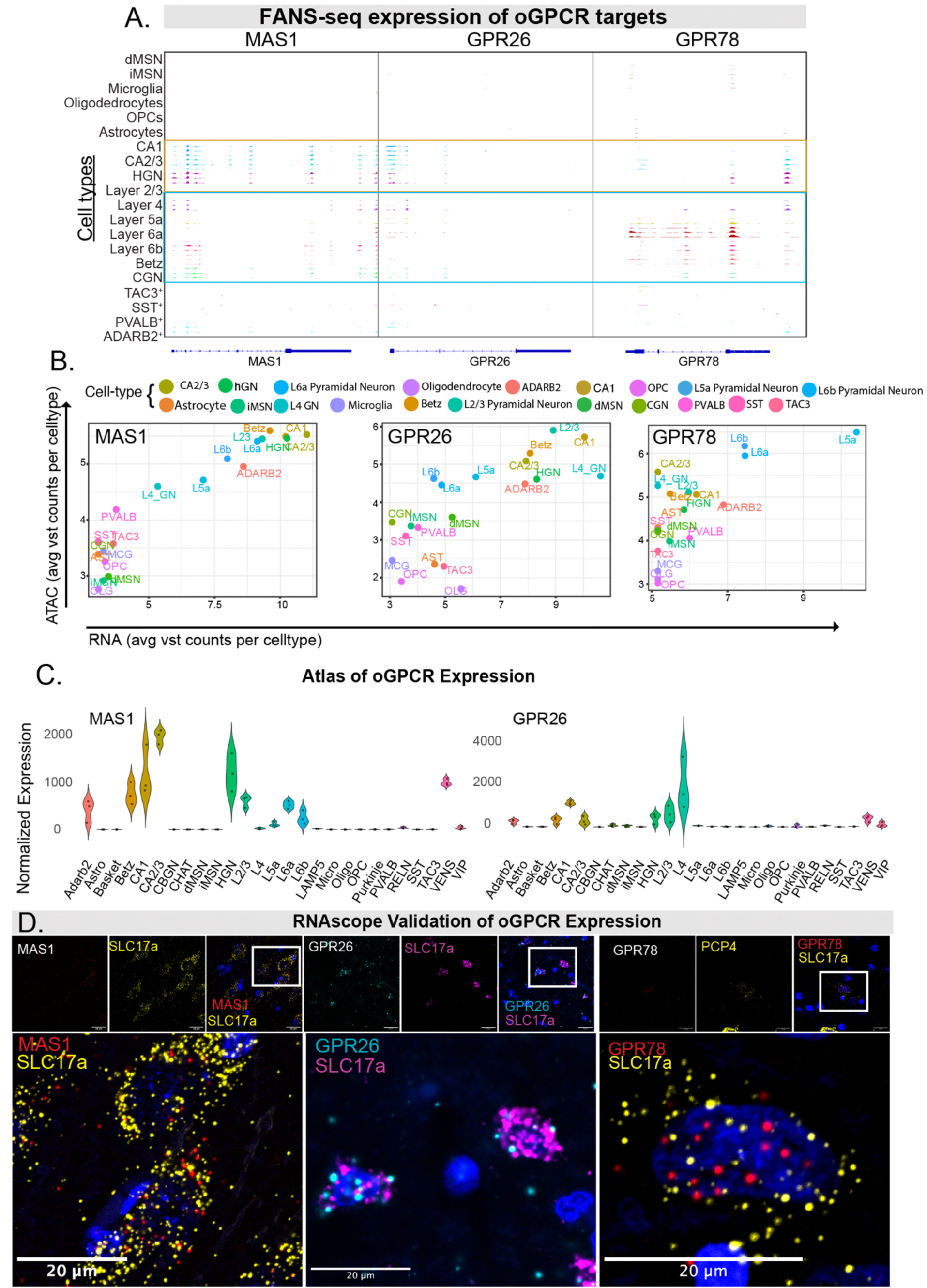
Cell-type enrichment of oGPCRs within the human neocortex and hippocampal formation. A. IGV plot of ATAC-accessibility for 3 enriched oGPCRs across 27 cell types (hippocampal cell types outlined in yellow, neocortex outlined in blue). B. Correlation analysis between ATAC-accessibility and RNA expression of each oGPCR displaying enrichment in ENs. C. Web-atlas RNA expression of *MAS1* and *GPR26* showing selective enrichment in ENs of the human brain. D. RNAscope validation of oGPCR cell-type enrichment of MAS1 in the CA1 region of the hippocampus (left) overlayed with EN marker SLC17a7, *GPR26* in the cortex (BA9) overlayed with EN marker SLC17A7 (middle) and GPR78 in cortical L5 overlayed with deep-layer marker *PCP4*.

In addition to the major EN cell-types of the cortex, FANS-seq allowed us to look for expression of oGPCR targets in rare cell populations such as Betz cells and Von Economo neurons of the cortex. Mapping oGPCR targets for potential selective neuromodulation of these rare populations is crucial in diseases such as amyotrophic lateral sclerosis (ALS) that display cell-type specific degeneration of Betz cells and in AD where the preservation of Von Economo neurons have been implicated in cognitive resilience to disease (*51, 52*). To assess if any of the cortical cell-specific oGPCRs displayed expression in these populations we first screened the FANS-Seq expression data within these sorted populations and observed expression of both *GPR26* and *MAS1* in both Betz cells and VENs cells (Fig. S5A). We validated the expression of both oGPCRs by looking for co-labeling with a previously identified Betz and VENS cell marker POU Class 3 Homeobox 1 (*POU3F1*) (Fig. S5B). Additionally, we observed selective enrichment of *GPR63* within *PCP4* positive Purkinje cells of the human cerebellum (Fig. S5C, D).

Through screening the expression of oGPCRs in neurons across the human brain, we observed one receptor with high enrichment in Purkinje cells of the cerebellum, *GPR63* (Fig. 1D, Fig. S6A). While this receptor showed many-folds enrichment in human Purkinje cells and conserved enrichment in these cells of the mouse brain (Fig. S6B), analyses of both FANS-seq and ATAC-seq data revealed that this receptor’s gene is both accessible and transcribed in many EN populations of the cortex and hippocampus, as well as in direct- and indirect-pathway MSNs (Fig. S6C). To confirm these results, we performed RNAscope on human caudate putamen and observed localized expression of *GPR63* transcripts within DARPP32 positive MSNs (Fig. S6D). Thus, although this receptor is highly enriched in certain cell-types such as Purkinje cells, molecular profiling reveals a broader expression pattern that evolutionary diverges from the selective expression observed in Purkinje cells of mice.

### oGPCR expression across human brain interneuron populations

Throughout the brain, the integration of both excitatory and inhibitory signaling is governed by a diverse set of interneurons regulating both local microcircuits and brain-wide functionality (*53*). Altered synaptic function of interneurons has been implicated in neurodevelopment, neuropsychiatric, and neurodegenerative conditions (*54–56*). Thus, defining cell-type specific ways to pharmacologically regulate interneuron function is essential to many neurological diseases. Here, we dissected numerous interneuron populations across the human striatum and neocortex to define enriched expression of oGPCRs for circuit modulation. Across all oGPCRs screened, only *GPR83* showed selective enrichment in interneuron subtypes (Fig. 1D). Enriched GPR83 RNA expression was observed across different interneuron populations of the striatum including PVALB+, SST+, TAC3+, and CHAT+ cells of the human striatum (Fig 6A). This enriched expression was confirmed by ATAC-accessible peaks within the *GPR83* gene in PVALB+, SST+, and TAC3+ interneurons (Fig. 6B). Interestingly. *GPR83* expression and accessibility was also observed in granule cells of the cerebellum and hippocampus, as well as, in EN populations of the hippocampus, but to a far lesser degree than striatal interneurons (Fig. 6B). To confirm *GPR83* enrichment in striatal interneuron subtypes we performed RNAscope for *GPR83* with interneuron markers SST and PVALB and we observed highly selective expression of *GPR83* in both SST and PVALB interneuron subtypes (Fig. 6C). These data demonstrate *GPR83* as a potential GPCR for selective modulation of interneuron activity.

**Figure 6.**
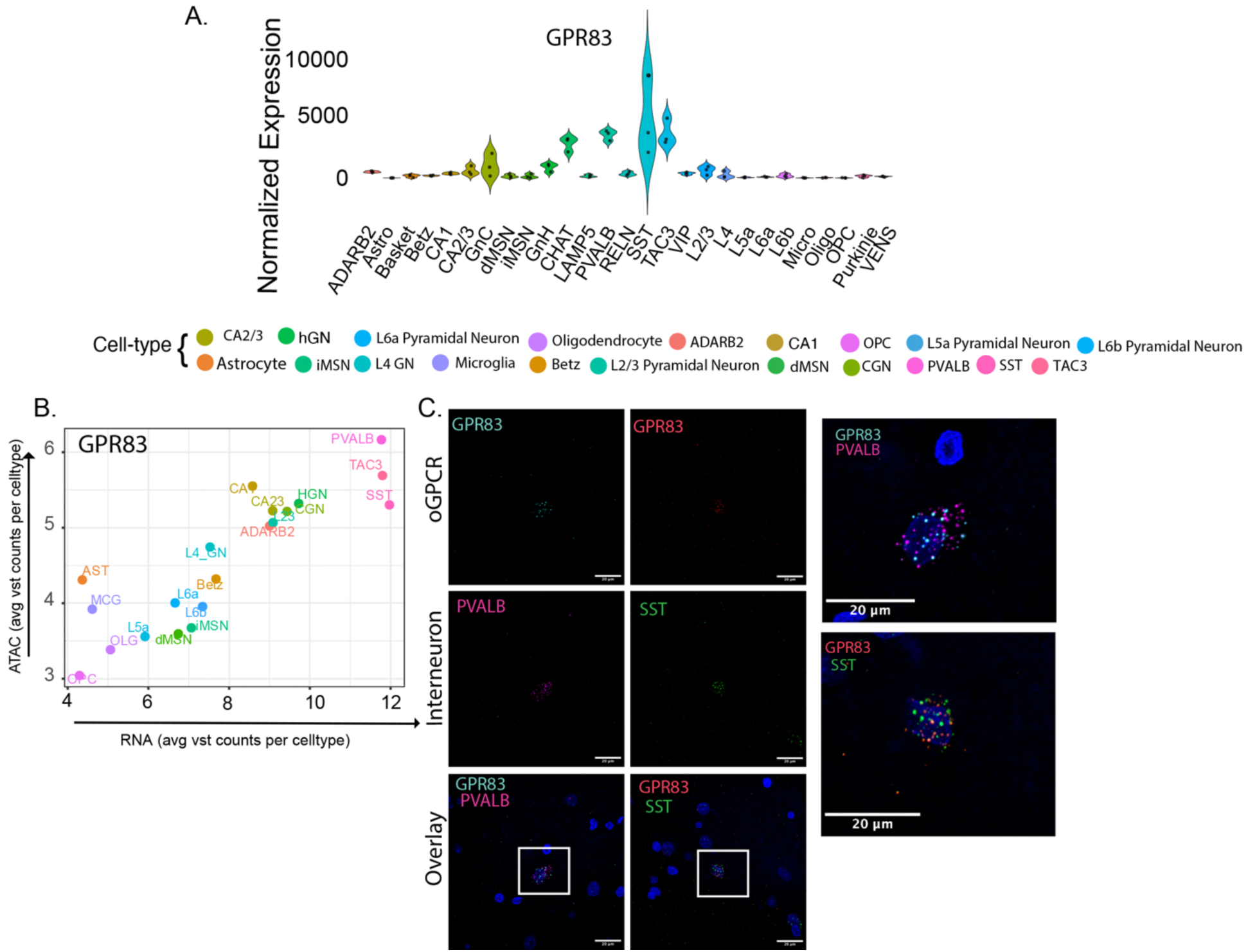
Enriched expression of GPR83 in specific interneuron subtypes. A. Normalized number of FANS-seq reads mapped to *GPR83* in human brain cell types analyzed. B. ATAC-mRNA correlations of *GPR83* accessibility and expression across human cell types. C. RNAscope-based detection of *GPR83* transcripts on sections of human putamen, overlayed with the signal from transcripts of interneuron subtype-specific marker genes *SST* and *PVALB*.

## DISCUSSION

Here, we have built an atlas of oGPCR expression across multiple regions of the human brain, identifying several GPCRs with cell-type specific expression. The characterization of each of these receptors is crucial to both deorphanizing efforts, and to generate interest in these receptors as potential targets for modulation of specific CNS circuits in the human brain in neurological and neurodegenerative disease. An example of this approach is the development of GPR6 inverse agonists as a non-dopaminergic and more selective therapy for Parkinson’s disease as a result of the selective expression of GPR6 in the indirect striatopallidal pathway (8,57). Promising Phase II clinical studies have resulted in the advance of these drugs into Phase III trials for treatment of both motor and non-motor disturbances occurring in PD (8,57). Furthermore, agonists targeting the MSN-specific receptor GPR52 are being evaluated as potential therapies for schizophrenia and psychosis (*57*), and preclinical studies of GPR88 suggests that it may be a useful target to address alcohol-use disorders (*58, 59*).

Understanding oGPCRs cell-type specific functional contributions to behavior may ultimately lead to de-orphanization of these receptors. For example, recent studies in mice have implicated an endogenous role for GPR139 in noncanonical opioid signaling with agonism observed via dynorphins in the medial habenula (*60*). Cells of the human striatum are heavily integrated with opioid circuits; thus GPR139 expression in human MSNs (described here) likely mediates a similar mechanism of dynorphin regulation (*61*). Here, we also have demonstrated highly selective expression of GPR101 in both dMSNs and iMSNs. Human clinical reports indicated that duplication of GPR101 leads to the developmental disorder X-linked acrogigantism (X-LAG)(*62*). Whereas studies in mice have shown hypothalamic GPR101 functions related to growth hormone signaling, it remains unclear how GPR101 expression in MSNs contributes to this disease or whether growth hormone signaling through GPR101 may create cross-talk within human CNS dopaminergic circuitry (*63, 64*). Overall, this atlas helps define targeting strategies for receptors such as GPR139 and GPR101 and highlights expression patterns that may mediate previously unknown biological functions. This atlas may also help resolve off-target effects when these receptors are pharmacologically manipulated.

Non-neuronal cell-types such as microglia and Oligos/OPCs displayed a high number of cell-type enriched oGPCRs. We observed a microglia-specific signature for the proton sensing GPCR, GPR65, which has well documented roles in peripheral inflammation, macrophage activation and tumorigenesis(*65, 66*). Previous studies have also implicated GPR34 and GPR141 in macrophage and microglial regulation in both the periphery and the CNS(*67, 68*). We further detected cell-type enrichment of GPR160 within microglia. Recent work has debated whether the endogenous ligand for GPR160 is cocaine- and amphetamine-regulated transcript peptide (CARTp), with some studies suggesting that GPR160 mediates this ligand’s role in pain (*69, 70*). Conversely, other *in vitro* studies fail to observe CARTp binding to GPR160(*71*). Here, we propose a translational approach to resolve GPR160 functionality by studying GPR160’s actions in microglial-specific cell lines and assays. FANS-based segmentation of cell-types allowed us to define Oligo/OPC specific receptors such as GPR37, which is oligodendrocyte specific, and GPR17, which displays much higher enrichment in OPCs. While both receptors have been implicated in myelination (*72, 73*), their translational targeting highlights lineage-specific modulation of mature or immature cell populations. Of note, we also observed shared expression of GPR37L1 in human astrocytes as well as oligodendrocytes. Recent studies implicate an astrocytic role of GPR37L1 in neuropathic pain in mice (*45*) and delineating oligodendrocyte roles for this receptor represents another future translational direction.

The only major cell-class without cell-type specific oGPCR expression was ENs of the neocortex and hippocampus. Although we identified neuronally enriched receptors in these brain regions, no receptor defined a specific cortical layer or any subregion of the hippocampal formation. However, expression of certain receptors did distinguish deep versus superficial neocortical layers, as shown by GPR78’s high enrichment in deep layers 5 and 6 of the neocortex. Importantly, FANS-seq allowed us to profile the orphan receptor landscape in rare cortical populations such as Betz and VENs cells, in which molecular characterization of modestly expressed drug targets is challenging using other sequencing modalities.

Importantly, while we demonstrate here that several oGPCRs are enriched in selective cell-types, some oGPCRs expressed in inhibitory MSNs are also expressed in inhibitory interneuron subtypes of the striatum (GPR149 and GPR88), albeit at lower levels. Furthermore, oGPCRs expression patterns observed in mice are not always evolutionally conserved in humans. For instance, studies in mice report GPR63 expression in Purkinje cells of the cerebellum and pyramidal cells of the hippocampus (*74*). In contrast, we find that while GPR63 is heavily enriched in Purkinje cells, it is also expressed broadly throughout many layers of the human neocortex and in both indirect and direct projecting MSNs of the striatum. In these cases it is important to delineate which cell-types may be driving particular phenotypes that appear mediated by these receptors, if the expression is precisely conserved in preclinical model systems, and how altered expression may influence therapeutic targeting of each receptor.

An interesting aspect of GPCR expression highlighted here is the overall biological redundancy of GPCRs within select cell-types of the CNS. While some biological segmentation of receptors in circuits is well characterized (DRD1 excites direct pathway MSNs, DRD2 inhibits indirect MSNs), it remains largely unknown why MSNs and other cell-types express multiple Gs-coupled GPCRs (GPR101, GPR88, GPR52, GPR6). This could reflect differential input and ligand specificity within these cell types such that each oGPCR has a unique circuit-specific action stimulated by its endogenous ligand either separately or in combinations with other ligands. Conversely, given the high constitutive activity of many oGPCRs, it’s possible that their main function is to set the baseline activity of critical signaling pathways. In that case, transcriptional and epigenetic mechanisms, as well as regulation of their subcellular localization, may be fundamental for their physiological roles in these circuits. Finally, the activity of these receptors may be mediated through protein-protein interactions via homo-dimerization or hetero-dimerization with other receptors in these cell-types (*75*). We conclude that further studies assessing each receptor’s structure, subcellular localization, downstream signaling pathways, transcriptional regulation, and pharmacogenomic responses will shed light on each individual receptor’s role in the brain. Nevertheless, definition of oGPCR expression patterns in the human brain is a first step towards understanding how these receptors function within specific cell-types and circuits, their potential signaling roles, and their implications in neurological disease.

## LIMITATIONS OF THE STUDY

Here, we utilized FANS-seq data to define cell-type specific oGPCRs throughout the human brain and created a resource of expression patterns for translational targeting. Importantly, our high-depth FANS-seq approach enabled the detection of oGPCRs that are undetectable with current methods, providing the first comprehensive view of cell-type–specific oGPCR expression in the human brain. However, future spatial and single-cell technologies with higher sensitivity and resolution may reveal additional oGPCR diversity that is not captured in our dataset. While this atlas serves as a first step in understanding the expression and function of these receptors, our analysis was constrained to four brain regions, so the atlas may not fully capture oGPCR diversity across the entire human CNS. Furthermore, these characterizations are based on mRNA detection and chromatin accessibility and thus lack information about proteomic regulation of each receptor, including activity-, internalization-, or disease-dependent regulation of oGPCRs that may alter expression patterns. Finally, while the expression patterns observed here highlight potential ways oGPCRs may be targeted in select cell-types of the human brain, ultimately functional studies in experimental model systems and clinical settings will be required to determine whether modulating the activity of these receptors can safely and effectively translate to therapeutics in neurological diseases.

## MATERIALS AND METHODS

### Study Design

Here, we utilized datasets of fluorescence activated nuclear sorted (FANS) and deeply profiled nuclei from human post-mortem brains to generate an atlas of orphan G-protein coupled receptors (oGPCR) expression across multiple regions of the human brain. We identified 22 oGPCRs that displayed cell-type RNA enrichment patterns and confirmed the cell-type enrichment of each receptor via FANS-ATAC-seq chromatin accessibility. From sections of human donors, we validated the expression of these oGPCRs within specific cell types and generated an interactive open-source web application to serve as a neuro-resource.

#### Human Samples

Deidentified tissue samples analyzed in this study were determined to be exempt from Institutional Review Board review according to 45 CFR 46.102 (f). For this work, fresh frozen brain samples were obtained from Miami’s Brain Endowment Bank, University of Washington BioRepository and Integrated Neuropathology Laboratory, Columbia University Alzheimer’ Disease Research Center, University of Maryland, Science Care, and Netherlands Brain Bank or through the National Institutes of Health (NIH) NeuroBioBank and sourced from either the Harvard Brain Tissue Resource Center, The University of Michigan Brain Bank or the NIH Brain & Tissue Repository-California, Human Brain & Spinal Fluid Resource Center, VA West LA Medical Center (Los Angeles, CA). Drug addiction and schizophrenia as well as clinical evidence of brain cancers were reasons for sample exclusion, whereas samples from donors with a history of other non-brain cancers and diabetes were accepted. Caudate nucleus, putamen, BA4, hippocampus, and cerebellum were used for immunohistochemistry.

### Nuclei Isolation, sorting, and FANS-seq

Nuclei isolation, sorting, and FANS-seq was conducted as previously described(*17–20*). Briefly, Frozen human tissue samples were homogenized through dounce-homogenization as described (*28*), The homogenate was brought to 24% iodixanol concentration and layered on top of a 27% iodixanol and Centrifugation was performed for 30 minutes at 4°C, 10,000 RPM Pelleted nuclei were resuspended in homogenization buffer. nuclei were fixed by adding 1mL of Homogenization buffer with 1mM EDTA and 1% formaldehyde (PFA) for 8 minutes at room temperature. The reaction was quenched through the addition of 0.125 M glycine. After 5 minutes of incubation, nuclei were washed once in homogenization buffer, followed by a transfer wash into blocking buffer containing Triton 0.05% X-100. Nuclei were blocked and permeabilized in Triton X-100 containing blocking buffer for 30 minutes at RT. After isolation, nuclei were resuspended in resuspension buffer in 100μl total volume and incubated with primary antibodies, secondary antibodies were added to the nuclei at a dilution of 1:1000, for a 30-minute incubation at RT. Three washes in wash buffer were followed by the last resuspension of nuclei in a DAPI containing wash buffer solution. Nuclei were separated by FACS immediately or stored overnight (ON) at 4°C. For neocortex samples sorting and gating strategies were the same as previously described (*19*), for striatum and cerebellum sorting and gating strategies were the same as previously described (*17, 18*), and hippocampal sorting and gating strategies were performed as described (*17*). For RNAseq experiments, RNA was extracted from fixed and sorted nuclei using the Qiagen AllPrep DNA/RNA FFPE kit. DNA from all samples was isolated and purified Genomic DNA was purified using AllPrep DNA/RNA FFPE Kit. Libraries were assessed for concentration and quality using the Qubit and Tapestation assays. Samples were pooled and sequenced using the NovaSeq SP 2×150 with the goal to reach a read depth of approximately 40 million reads per sample.

#### Data Processing

RNA and ATAC datasets were processed consistently with previous publications (*17–19*). Briefly, for all RNAseq data, sequence and transcript coordinates for human hg38 UCSC genome and gene models were retrieved from the BSgenome.Hsapiens.UCSC.hg38 and TxDb.Haspiens.UCSC.hg38 Gene Bioconductor libraries. Genomic alignment of RNAseq reads were aligned to the genome using Rsubread’s subjunc method (*76*). Reads in genes were counted using the featurecounts function (*77*) within the Rsubread package against the full gene bodies for nuclear RNAseq (*78*). Gene-level count matrices were imported into DESeq2 (*79, 80*) and used to construct a DESeq dataset with cell type specified as the design factor. Size-factor normalization was performed using the DESeq2 median-of-ratios method to account for differences in sequencing depth across samples. Normalized counts were extracted and used for downstream analyses. For visualization and exploratory analyses, normalized counts were log2-transformed as log2 (count +1). Gene identifiers were mapped from Entrez IDs to gene symbols using org.Hs.eg.db, and genes without mapped symbols were excluded.

For ATAC-seq data, reads were aligned with the human hg38 genome from the BSgenome.Hsapiens.UCSC.hg38 Bioconductor package (version 1.4.1) fragments between 1 and 5,000 base pairs long were considered correctly paired. MACS2 software in BAMPE mode was used for peak calling (*81, 82*). Differential ATACseq signals were identified with the DESeq2 package (*79, 80*) (version 1.20.0) and high confidence consensus peaks were annotated to TSS based on proximity using the ChIPseeker package (*83*) (version 1.28.3).

For ATAC-RNA correlated expression, ATACseq and RNAseq peak to gene linkages and correlations were performed as previously described (*17*). Briefly, normalized peak and gene counts are generated using RNAseq and ATACseq datasets from the same donor and region. For every gene, accessible regions within 500kb of the TSS are tested for significant correlations with the expression of that gene and assigned a correlation coefficient, p-value and FDR. Significant Peak to gene linkages were only considered for peaks with an FDR < 0.05 and correlation coefficient greater than 0.3 or less than −0.3.

#### Sample Preparation

Formalin-fixed, paraffin embedded (FFPE) samples from desired brain regions were provided by the brain banks and shipped at room temperature to Rockefeller University. FFPE samples were sectioned on a Leica HistoCore AUTOCUT Rotary Microtome (5µM). Samples were mounted on SuperFrost Plus slides and were stored at room temperature. Prior to labeling, sections were subjected to a high intensity photobleaching protocol(*84*) (48-72hr)

#### RNAscope and Immunohistochemistry

In situ Hybridization and immunofluorescence experiments were performed over the course of this work using RNAscope Multiplex Fluorescence V2 Assay (Advanced Cell Diagnostics, ACD was performed according to the manufacturers protocol with minor modifications on formalin-fixed, paraffin embedded (FFPE) tissue samples. Briefly, FFPE sections were baked at 60°C for 1 hour, followed by deparaffination in fresh xylene 2 times 5 minutes each and dehydrated in 100% ethanol 2 times for 2 minutes each. Sections were air dried and treated with RNAscope hydrogen peroxide for 10 minutes at room temperature. Antigen retrieval was carried out by immersing slides in RNAscope Target Retrieval Buffer at 100°C for 15 minutes, followed by rinsing in distilled water and 100% ethanol. Tissue was permeabilized with RNAscope Protease Plus Reagent for 30 minutes and sections were then hybridized with RNAscope probes targeting genes of interest for 2 hours at 40°C. Sections were then put through a series of amplification steps, and fluorophores were assigned using TSA-conjugated Alexa Flour dyes. Between each amplification step slides were washed with RNAscope Wash Buffer. Following the completion of RNAscope signal development, sections were blocked in 3% Donkey serum with .3% Triton X-100 for 1 hour at room temperature. Primary antibodies against cell-type-specific markers (Iba1, Darpp32, GFAP 1/500) were applied and incubated overnight at 4°C in blocking buffer. Sections were washed in PBS and incubated with species-appropriate secondary antibodies conjugated to Alexa Fluor 488 (1/500). Nuclei were counterstained with RNAscope DAPI and cover slipped using Prolong Gold Antifade reagent. Imaging of fluorescent samples was performed on the LSM710 Zeiss Confocal Microscope, using Zeiss ZEN microscopy software. ISH images were processed using Fiji.

#### GPCRxplorer

GPCRxplorer is a custom interactive web application developed in R to visualize RNA-seq derived gene expression across defined neural and glial cell types. The application was built using the Shiny framework in Rstudio. Preprocessed RNA-seq data were loaded into R as a serialized object containing normalized gene expression values, annotated by gene symbols and cell types. Class a orphan GPCRs were made available for selection in the user interface. The application generates violin plots depicting expression distributions across cell types, displayed either on a linear scale or log-transformed scale. Individual sequencing experiments were overlaid in violins to display variability. Plot rendering and layout were implemented using ggplot2 and patchwork R packages. The application included the function to export figures as scalable vector graphics (SVG) files using the svglite package. GPCRxplorer provides a platform for exploratory and comparative analysis of GPCR expression patterns across brain cell types using RNA-seq data.

## List of Supplementary Materials

Fig. S1 to S6

Data File S1 to S4

## Acknowledgments

We would like to acknowledge all members of the Heintz laboratory for thoughtful discussions pertaining to this manuscript. We also acknowledge the Defense Health Agency Neuroanatomical Collections Division of the National Museum of Health and Medicine for permitted use of the image in Figure 1A.

## Funding

This work was supported by funds from the Fisher Center for Alzheimer’s Research (NH).

## Author contributions

Conceptualization: AU, PW, IIT, NH. Methodology: AU, PW, CP, MB, ES, KM. Investigation: PW, AU, FF, MB. Visualization: CP, MB, AU, PW. Funding acquisition: NH. Project administration: IIT, NH. Supervision: AU, IIT, NH. Writing – original draft: AU, PW. Writing – review & editing: AU, PW, IIT, NH.

## Competing interests

NH is a consultant for Cerevance

## Data and materials availability

All data will be made available upon request.

## Supplementary Materials

**Supplementary Figure 1.**
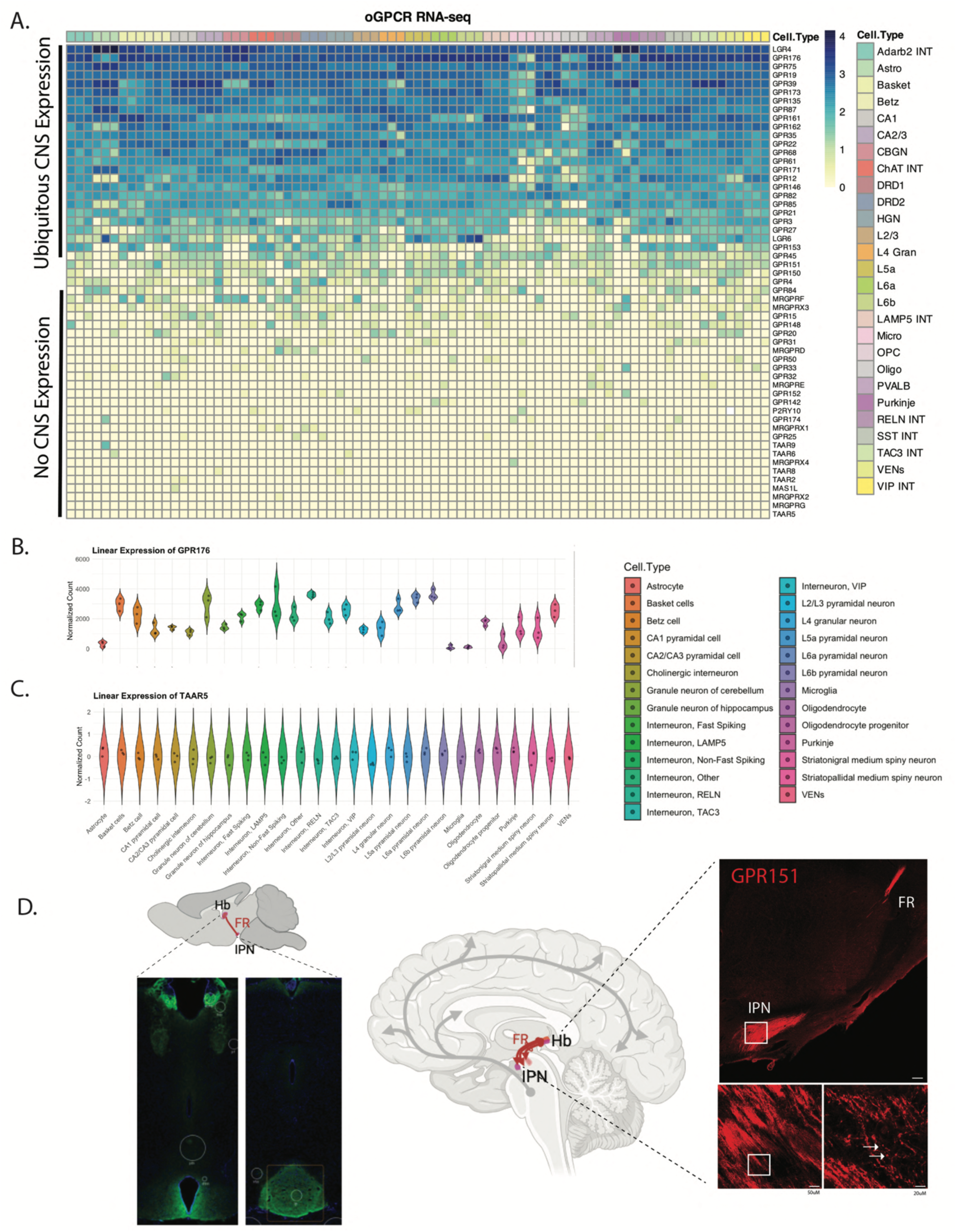
Non-selective or undetectable expression of oGPCRs in human brain cell types analyzed. A. Heatmap of log gene counts showing FANS-seq expression of oGPCRs across human brain regions assessed. B. Web-atlas violin plots displaying ubiquitous expression of GPR176 throughout the human brain. C. Web-atlas violin plots displaying undetectable expression of TAAR5 across the human brain regions assessed. D. Conserved cell-type expression of GPR151 in the mouse (left, The Human Protein Atlas) and in the human habenula (Hb)-fasciculus retroflexus (FR)-interpeduncular nucleus (IPN) tract (right).

**Supplementary Figure 2.**
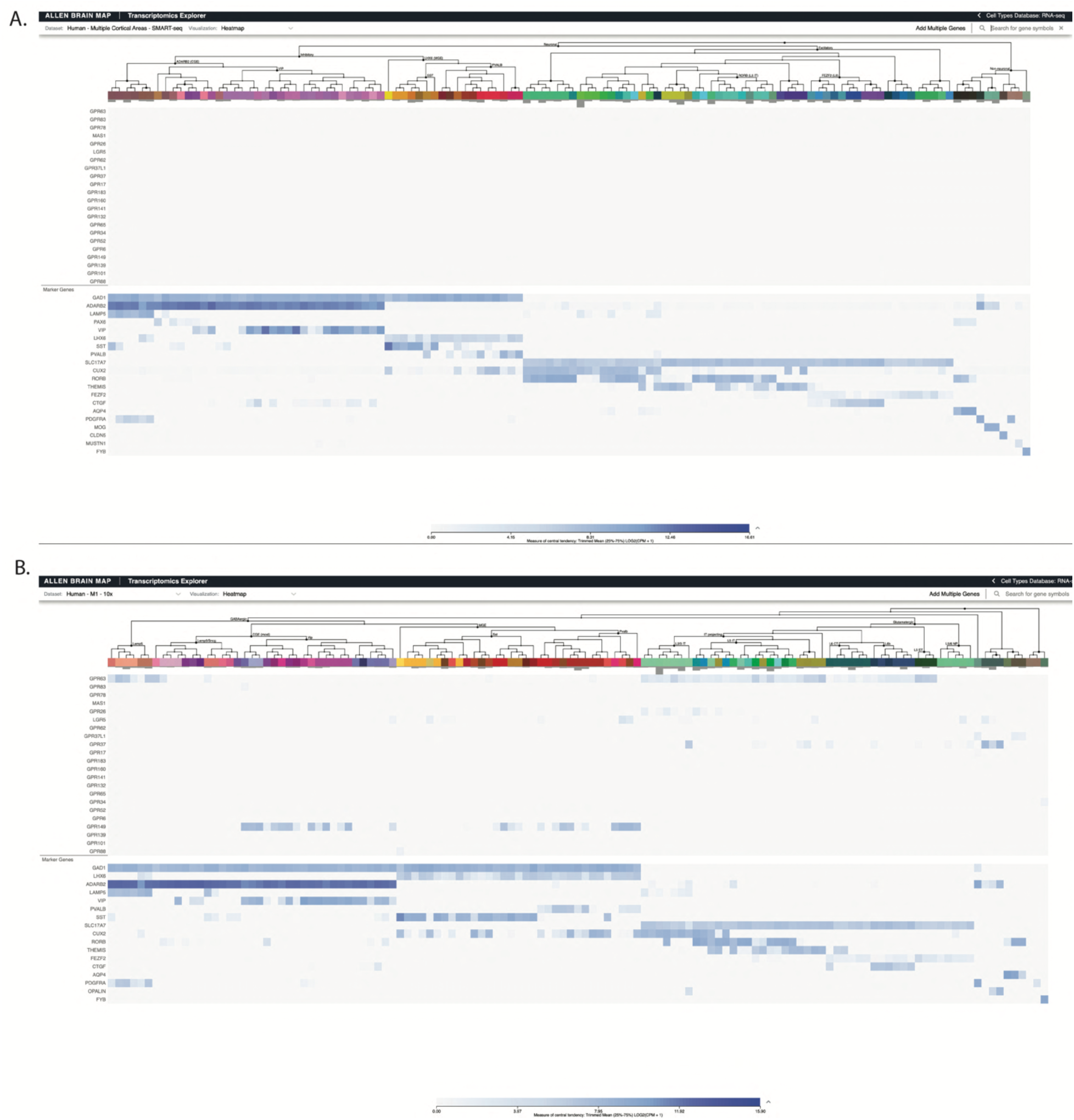
Available single-cell cell-type atlases for oGPCR expression from the Allen Brain Atlas (https://brain-map.org/our-research/cell-types-taxonomies/cell-types-database-rna-seq-data). A. Allen-Brain Atlas SMART-seq data from human neocortex searching for oGPCR expression. B. Allen-Brain Atlas 10x genomics single-cell atlas searching for oGPCR expression.

**Supplemental Figure 3.**
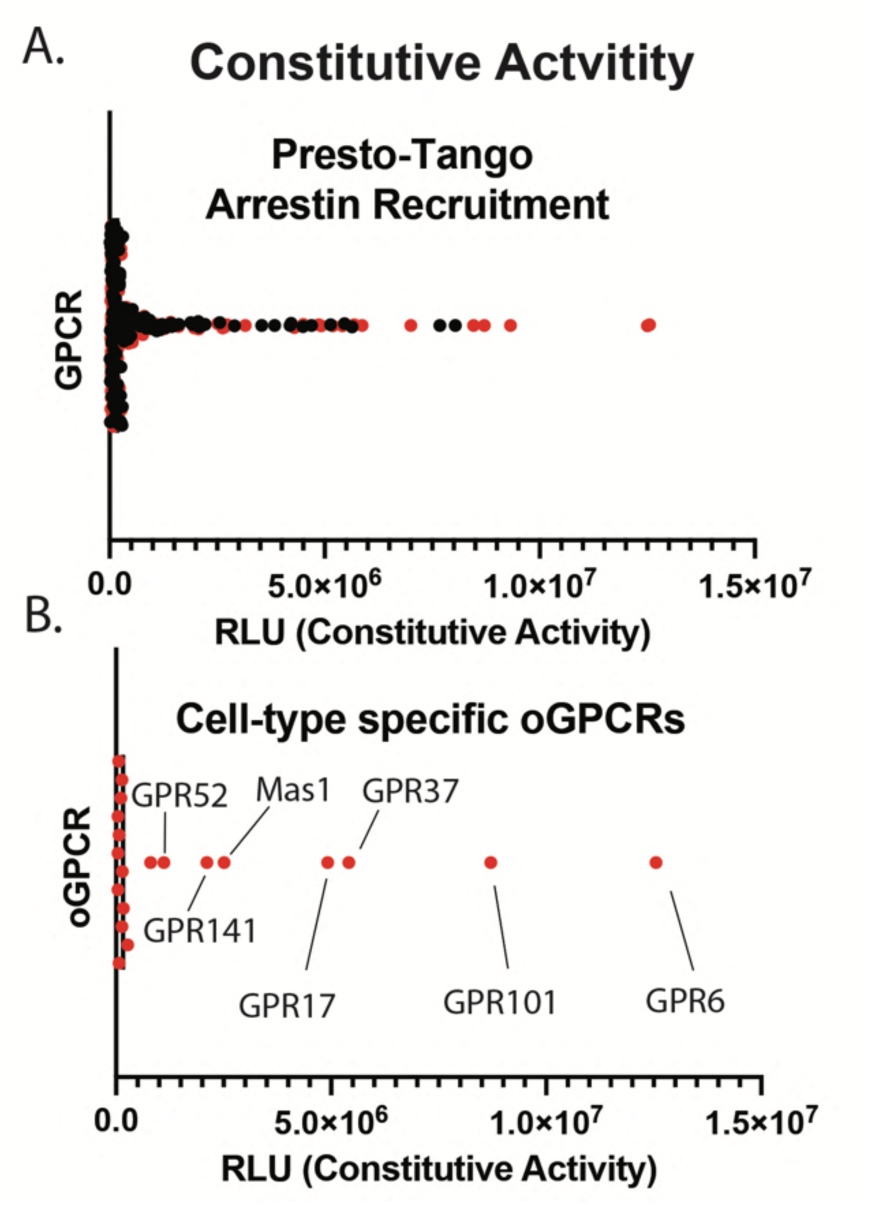
Constitutive activity of GPCRs. A. Presto-TANGO β-arrestin recruitment of all GPCRs tested from(*31*) oGPCRs highlighted in red. B. Isolated constitutive activity of cell-type specific oGPCRs assessed in this study.

**Supplemental Figure 4.**
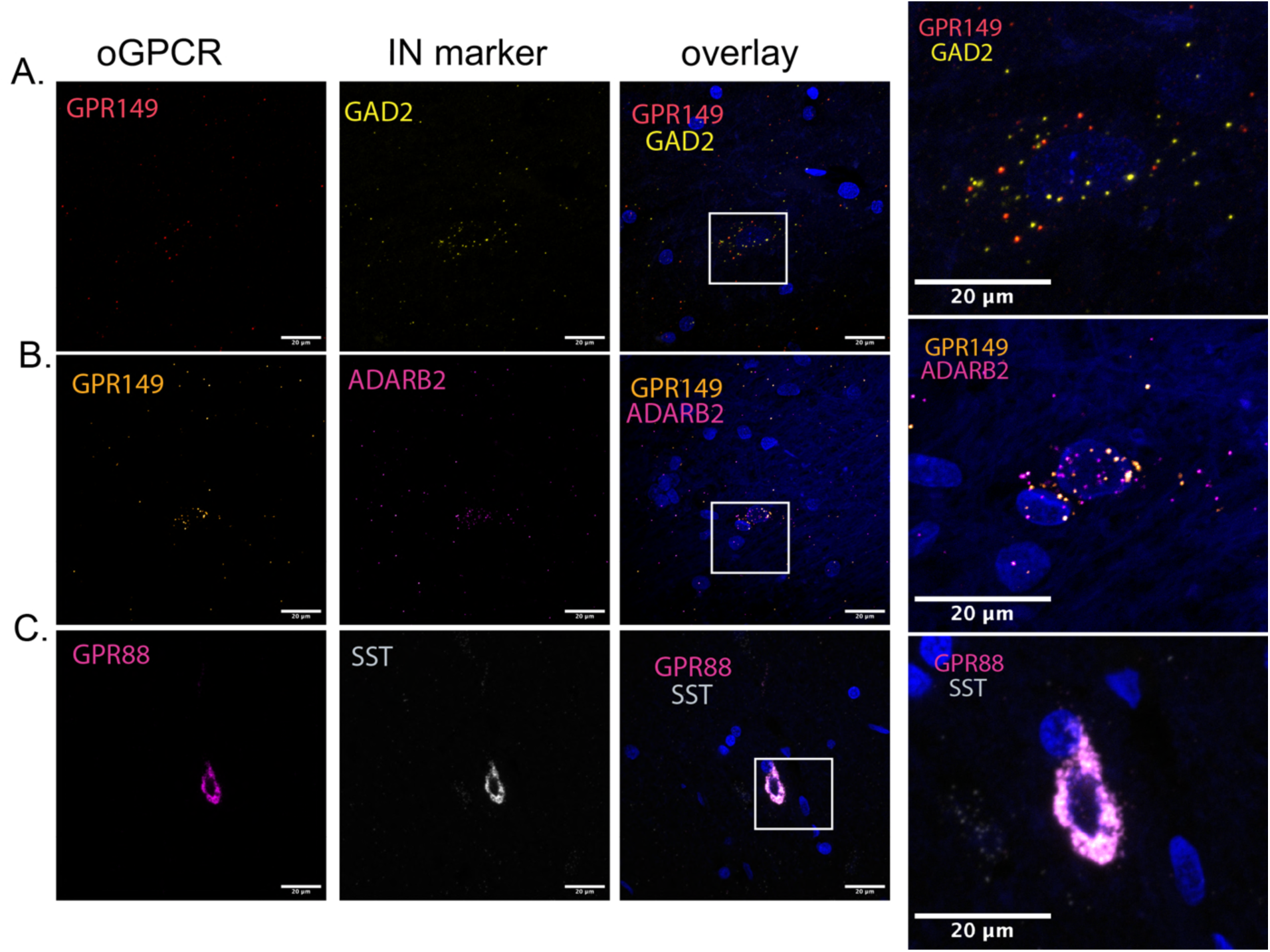
Detection of low expression levels of striatal enriched oGPCRs in interneurons. A. RNAscope of GPR149 overlayed with GABAergic marker GAD2 in human hippocampus. B. RNAscope of GPR149 overlayed with interneuron marker ADARB2 in human hippocampus. C. RNAscope of GPR88 overlayed with interneuron marker SST in human caudate.

**Supplementary Figure 5.**
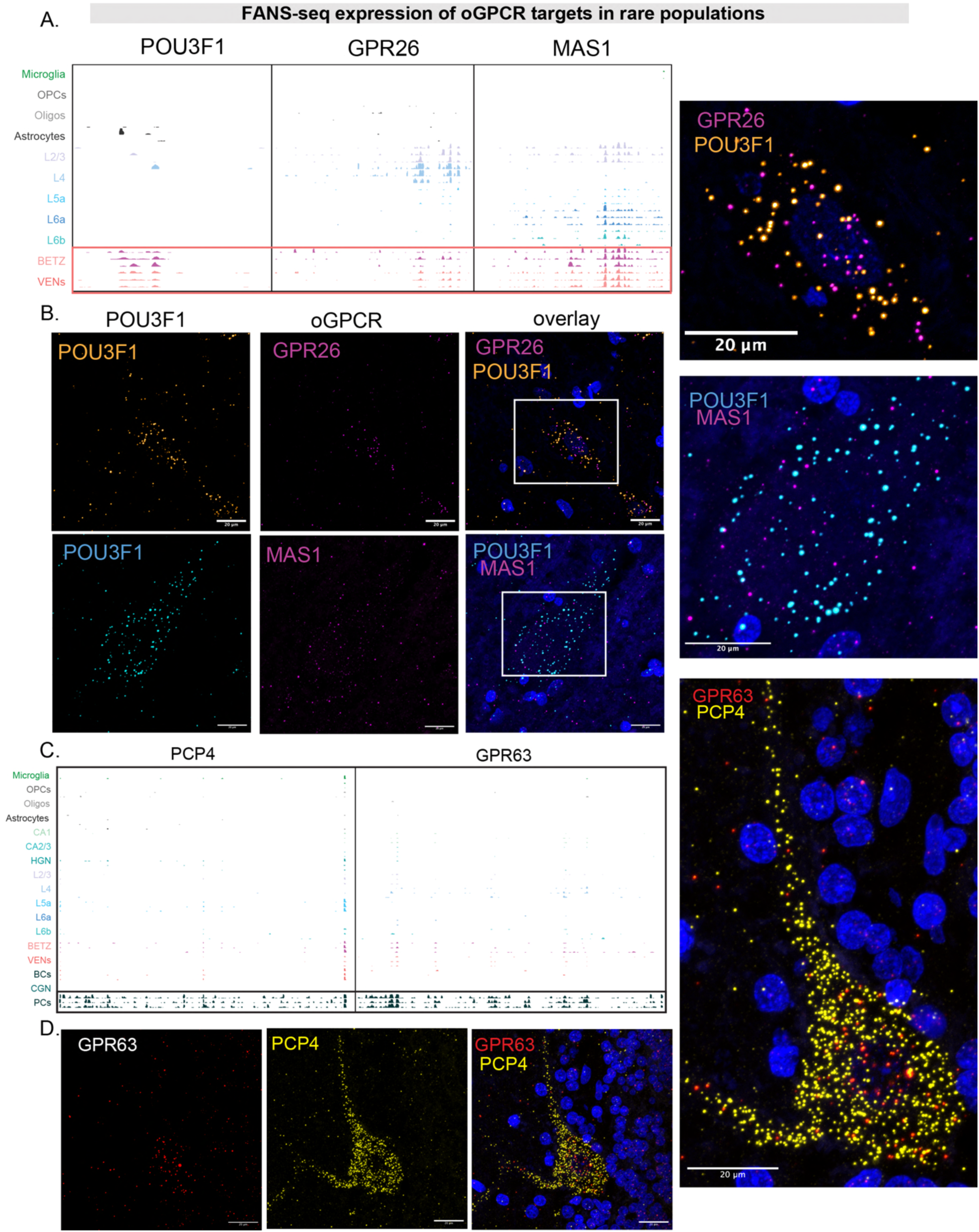
FANS-seq-based identification and RNAScope-based validation of oGPCRs expression in rare cell populations. A. IGV-based visualization of the distribution of FANS-seq reads over *GPR26, MAS1* and in *POU3F1* genes in selected cell type-specific datasets from human neocortex. B. RNAscope validation of GPR26 and MAS1 within large POU3F1+ cells of the human cortex. C. IGV of FANS-seq expression of *GPR63* showing enrichment in Purkinje cells. D. RNAscope-based detection of *GPR63* transcripts on sections of human cerebellum, overlayed with the signal from the transcripts of Purkinje cell-specific marker gene *PCP4*.

**Supplementary Figure 6.**
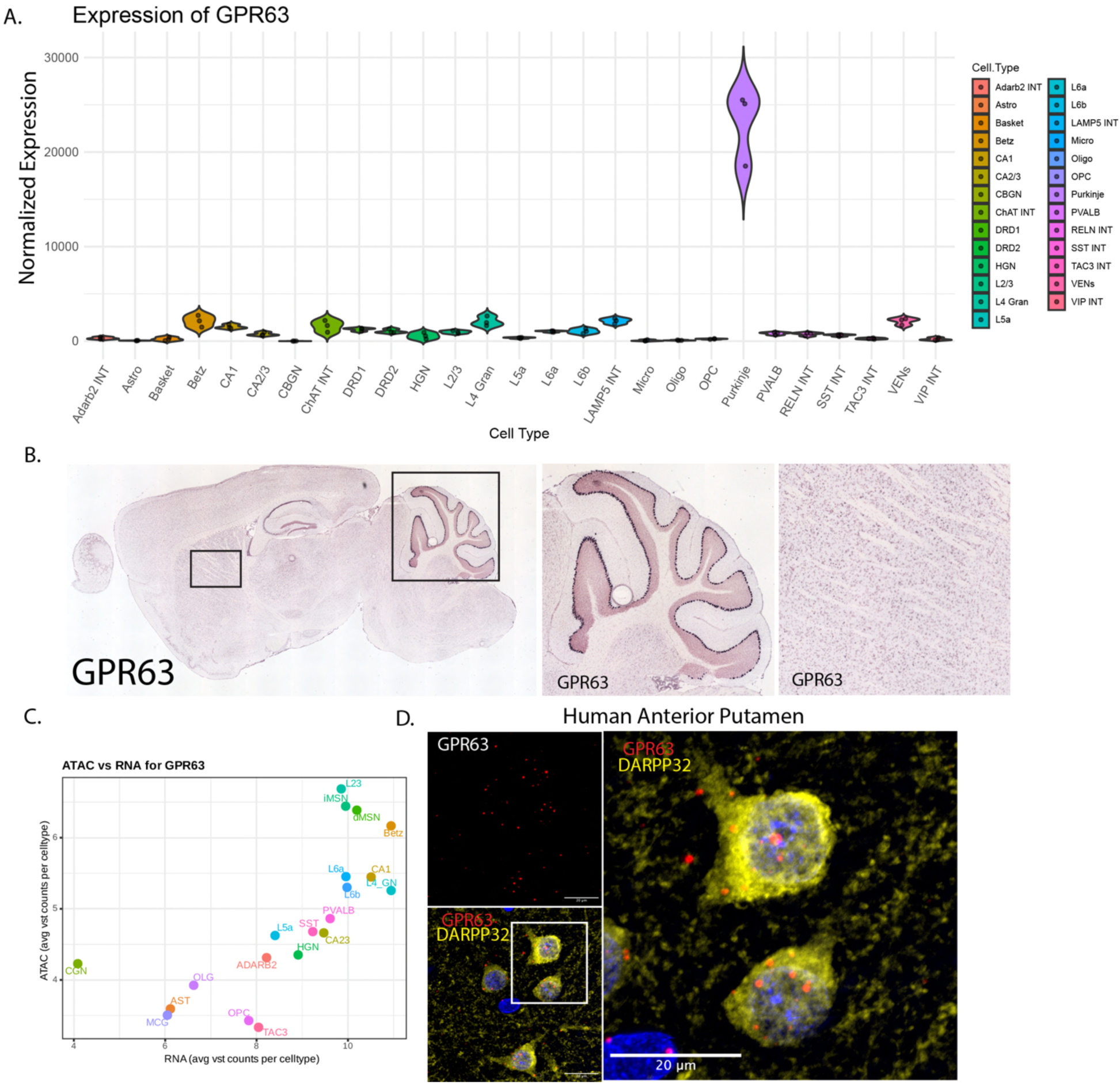
Evolutionary divergence and broader expression pattern of Purkinje cell enriched receptor GPR63 in human brain. A. Normalized number of FANS-seq reads mapped to *GPR63* in human brain cell types analyzed. B. Purkinje cell enrichment of GPR63 in mice cerebellum (Allen Brain ISH). C. RNA-ATAC correlation of GPR63 expression and accessibility across cell types in the human brain. D. RNAscope-based detection of *GPR63* transcripts on sections of human putamen overlayed with the signal from immunostaining of MSN marker DARPP32.

## Notes

https://cpressl.shinyapps.io/GPCRxplorer

### Competing Interest Statement

NH is a consultant at Cerevance.

## References and Notes

1. S. Liu, P. J. Anderson, S. Rajagopal, R. J. Lefkowitz, H. A. Rockman, G Protein-Coupled Receptors: A Century of Research and Discovery. Circ Res 135, 174–197 (2024).

2. J. S. Lorente et al., GPCR drug discovery: new agents, targets and indications. Nat Rev Drug Discov 24, 458–479 (2025).

3. T. S. Wong et al., G protein-coupled receptors in neurodegenerative diseases and psychiatric disorders. Signal Transduct Target Ther 8, 177 (2023).

4. I. Mantas, M. Saarinen, Z. D. Xu, P. Svenningsson, Update on GPCR-based targets for the development of novel antidepressants. Mol Psychiatry 27, 534–558 (2022).

5. K. Sriram, P. A. Insel, G Protein-Coupled Receptors as Targets for Approved Drugs: How Many Targets and How Many Drugs? Mol Pharmacol 93, 251–258 (2018).

6. A. T. Ehrlich et al., Expression map of 78 brain-expressed mouse orphan GPCRs provides a translational resource for neuropsychiatric research. Commun Biol 1, 102 (2018).

7. Y. Fang, T. Kenakin, C. Liu, Editorial: Orphan GPCRs As Emerging Drug Targets. Front Pharmacol 6, 295 (2015).

8. N. L. Brice et al., CVN424, a GPR6 inverse agonist, for Parkinson’s disease and motor fluctuations: a double-blind, randomized, phase 2 trial. EClinicalMedicine 77, 102882 (2024).

9. N. L. Brice et al., Development of CVN424: A Selective and Novel GPR6 Inverse Agonist Effective in Models of Parkinson Disease. J Pharmacol Exp Ther 377, 407–416 (2021).

10. D. H. Margolin, N. L. Brice, A. M. Davidson, K. L. Matthews, M. B. L. Carlton, A Phase I, First-in-Human, Healthy Volunteer Study to Investigate the Safety, Tolerability, and Pharmacokinetics of CVN424, a Novel G Protein-Coupled Receptor 6 Inverse Agonist for Parkinson’s Disease. J Pharmacol Exp Ther 381, 33–41 (2022).

11. H. Sun et al., First-Time Disclosure of CVN424, a Potent and Selective GPR6 Inverse Agonist for the Treatment of Parkinson’s Disease: Discovery, Pharmacological Validation, and Identification of a Clinical Candidate. J Med Chem 64, 9875–9890 (2021).

12. S. R. Benedetto et al., A Phase 2 Randomized Trial of NBI-1065846 (TAK-041) in Patients With Anhedonia Associated With Major Depressive Disorder: Results of the TERPSIS Study. J Clin Psychopharmacol 45, 432–440 (2025).

13. P. C. T. Hawkins et al., The GPR139 agonist TAK-041 produces time-dependent alterations to cerebral blood flow and reward system function in patients with schizophrenia: a randomised placebo-controlled trial. Psychopharmacology (Berl), (2025).

14. H. A. Reichard et al., Discovery of TAK-041: a Potent and Selective GPR139 Agonist Explored for the Treatment of Negative Symptoms Associated with Schizophrenia. J Med Chem 64, 11527–11542 (2021).

15. W. Yin et al., A phase 1 study to evaluate the safety, tolerability and pharmacokinetics of TAK-041 in healthy participants and patients with stable schizophrenia. Br J Clin Pharmacol 88, 3872–3882 (2022).

16. A. Bilal et al., A Randomized Controlled, Double-Masked, Crossover Study of a GPR119 Agonist on Glucagon Counterregulation During Hypoglycemia in Type 1 Diabetes. Diabetes 74, 1262–1272 (2025).

17. M. Baffuto et al., Epigenetic mechanisms governing cell type specific somatic expansion and toxicity in Huntington’s disease. bioRxiv, (2025).

18. K. Matlik et al., Cell-type-specific CAG repeat expansions and toxicity of mutant Huntingtin in human striatum and cerebellum. Nat Genet 56, 383–394 (2024).

19. C. Pressl et al., Selective vulnerability of layer 5a corticostriatal neurons in Huntington’s disease. Neuron 112, 924–941 e910 (2024).

20. C. Pressl, et al., Isolation and Molecular Profiling of Nuclei of Specific Neuronal Types from Human Cerebral Cortex and Striatum. Curr Protoc 4, e70067 (2024).

21. M. Doi et al., Gpr176 is a Gz-linked orphan G-protein-coupled receptor that sets the pace of circadian behaviour. Nat Commun 7, 10583 (2016).

22. J. Tang et al., GPR176 Promotes Cancer Progression by Interacting with G Protein GNAS to Restrain Cell Mitophagy in Colorectal Cancer. Adv Sci (Weinh) 10, e2205627 (2023).

23. J. Tian, Z. Huang, W. Zhang, Gpr176 modulates the firing pattern of parvalbumin-positive interneurons in the orbitofrontal cortex of mouse. Mol Brain 18, 81 (2025).

24. J. Broms, B. Antolin-Fontes, A. Tingstrom, I. Ibanez-Tallon, Conserved expression of the GPR151 receptor in habenular axonal projections of vertebrates. J Comp Neurol 523, 359–380 (2015).

25. Z. Yao et al., A high-resolution transcriptomic and spatial atlas of cell types in the whole mouse brain. Nature 624, 317–332 (2023).

26. X. Chen et al., A brain cell atlas integrating single-cell transcriptomes across human brain regions. Nat Med 30, 2679–2691 (2024).

27. K. L. Chiou et al., A single-cell multi-omic atlas spanning the adult rhesus macaque brain. Sci Adv 9, eadh1914 (2023).

28. X. Xu et al., Species and cell-type properties of classically defined human and rodent neurons and glia. Elife 7, (2018).

29. A. Jobe, R. Vijayan, Orphan G protein-coupled receptors: the ongoing search for a home. Front Pharmacol 15, 1349097 (2024).

30. A. L. Martin, M. A. Steurer, R. S. Aronstam, Constitutive Activity among Orphan Class-A G Protein Coupled Receptors. PLoS One 10, e0138463 (2015).

31. W. K. Kroeze et al., PRESTO-Tango as an open-source resource for interrogation of the druggable human GPCRome. Nat Struct Mol Biol 22, 362–369 (2015).

32. L. M. Yager, A. F. Garcia, A. M. Wunsch, S. M. Ferguson, The ins and outs of the striatum: role in drug addiction. Neuroscience 301, 529–541 (2015).

33. F. Plattner et al., The role of ventral striatal cAMP signaling in stress-induced behaviors. Nat Neurosci 18, 1094–1100 (2015).

34. Y. Han et al., Decoding the striatum of drug-naive patients with obsessive-compulsive disorder: a transcriptome and longitudinal functional magnetic resonance imaging study. Transl Psychiatry 15, 258 (2025).

35. S. Zhai, A. Tanimura, S. M. Graves, W. Shen, D. J. Surmeier, Striatal synapses, circuits, and Parkinson’s disease. Curr Opin Neurobiol 48, 9–16 (2018).

36. J. A. Girault, A. C. Nairn, DARPP-32 40 years later. Adv Pharmacol 90, 67–87 (2021).

37. C. Gao, J. Jiang, Y. Tan, S. Chen, Microglia in neurodegenerative diseases: mechanism and potential therapeutic targets. Signal Transduct Target Ther 8, 359 (2023).

38. O. Butovsky, H. L. Weiner, Microglial signatures and their role in health and disease. Nat Rev Neurosci 19, 622–635 (2018).

39. C. C. Hsiao et al., GPCRomics of Homeostatic and Disease-Associated Human Microglia. Front Immunol 12, 674189 (2021).

40. C. Gu et al., Role of G Protein-Coupled Receptors in Microglial Activation: Implication in Parkinson’s Disease. Front Aging Neurosci 13, 768156 (2021).

41. S. Hickman, S. Izzy, P. Sen, L. Morsett, J. El Khoury, Microglia in neurodegeneration. Nat Neurosci 21, 1359–1369 (2018).

42. D. Oz-Arslan, M. Yavuz, B. Kan, Exploring orphan GPCRs in neurodegenerative diseases. Front Pharmacol 15, 1394516 (2024).

43. K. E. Hopperton, D. Mohammad, M. O. Trepanier, V. Giuliano, R. P. Bazinet, Markers of microglia in post-mortem brain samples from patients with Alzheimer’s disease: a systematic review. Mol Psychiatry 23, 177–198 (2018).

44. J. An et al., G protein-coupled receptor GPR37-like 1 regulates adult oligodendrocyte generation. Dev Neurobiol 81, 975–984 (2021).

45. J. Xu, Z. Yan, S. Bang, D. Velmeshev, R. R. Ji, GPR37L1 identifies spinal cord astrocytes and protects neuropathic pain after nerve injury. Neuron 113, 1206–1222 e1206 (2025).

46. Q. Zhou, G. Choi, D. J. Anderson, The bHLH transcription factor Olig2 promotes oligodendrocyte differentiation in collaboration with Nkx2.2. Neuron 31, 791–807 (2001).

47. K. Khodosevich, C. M. Sellgren, Neurodevelopmental disorders-high-resolution rethinking of disease modeling. Mol Psychiatry 28, 34–43 (2023).

48. W. Jagust, Vulnerable neural systems and the borderland of brain aging and neurodegeneration. Neuron 77, 219–234 (2013).

49. J. Therriault et al., Staging of Alzheimer’s disease: past, present, and future perspectives. Trends Mol Med 28, 726–741 (2022).

50. H. Zeng et al., Large-scale cellular-resolution gene profiling in human neocortex reveals species-specific molecular signatures. Cell 149, 483–496 (2012).

51. H. Braak, A. C. Ludolph, M. Neumann, J. Ravits, K. Del Tredici, Pathological TDP-43 changes in Betz cells differ from those in bulbar and spinal alpha-motoneurons in sporadic amyotrophic lateral sclerosis. Acta Neuropathol 133, 79–90 (2017).

52. T. Gefen et al., Von Economo neurons of the anterior cingulate across the lifespan and in Alzheimer’s disease. Cortex 99, 69–77 (2018).

53. H. Markram et al., Interneurons of the neocortical inhibitory system. Nat Rev Neurosci 5, 793–807 (2004).

54. K. Ure et al., Restoration of Mecp2 expression in GABAergic neurons is sufficient to rescue multiple disease features in a mouse model of Rett syndrome. Elife 5, (2016).

55. Y. Arime et al., Activation of prefrontal parvalbumin interneurons ameliorates working memory deficit even under clinically comparable antipsychotic treatment in a mouse model of schizophrenia. Neuropsychopharmacology 49, 720–730 (2024).

56. S. Hijazi, A. B. Smit, R. E. van Kesteren, Fast-spiking parvalbumin-positive interneurons in brain physiology and Alzheimer’s disease. Mol Psychiatry 28, 4954–4967 (2023).

57. S. Ali, P. Wang, R. E. Murphy, J. A. Allen, J. Zhou, Orphan GPR52 as an emerging neurotherapeutic target. Drug Discov Today 29, 103922 (2024).

58. S. Ben Hamida et al., The GPR88 agonist RTI-13951-33 reduces alcohol drinking and seeking in mice. Addict Biol 27, e13227 (2022).

59. M. Fer et al., Discovery of BI-9508, a Brain-Penetrant GPR88-Receptor-Agonist Tool Compound for In Vivo Mouse Studies. J Med Chem 67, 11296–11325 (2024).

60. X. Li et al., Homeostatic scaling of dynorphin signaling by a non-canonical opioid receptor. Nat Commun 16, 6786 (2025).

61. G. Henriksen, F. Willoch, Imaging of opioid receptors in the central nervous system. Brain 131, 1171–1196 (2008).

62. C. Rodd et al., Somatic GPR101 Duplication Causing X-Linked Acrogigantism (XLAG)-Diagnosis and Management. J Clin Endocrinol Metab 101, 1927–1930 (2016).

63. Z. Yang et al., Structure of GPR101-Gs enables identification of ligands with rejuvenating potential. Nat Chem Biol 20, 484–492 (2024).

64. D. Abboud et al., GPR101 drives growth hormone hypersecretion and gigantism in mice via constitutive activation of G(s) and G(q/11). Nat Commun 11, 4752 (2020).

65. S. Bagchi, et al., The acid-sensing receptor GPR65 on tumor macrophages drives tumor growth in obesity. Sci Immunol 9, eadg6453 (2024).

66. I. Neale et al., Small-molecule probe for IBD risk variant GPR65 I231L alters cytokine signaling networks through positive allosteric modulation. Sci Adv 10, eadn2339 (2024).

67. T. Schoneberg, J. Meister, A. B. Knierim, A. Schulz, The G protein-coupled receptor GPR34 - The past 20 years of a grownup. Pharmacol Ther 189, 71–88 (2018).

68. A. Sawabe, S. Okazaki, A. Nakamura, R. Goitsuka, T. Kaifu, The orphan G protein-coupled receptor 141 expressed in myeloid cells functions as an inflammation suppressor. J Leukoc Biol 115, 935–945 (2024).

69. M. O. Job, M. J. Kuhar, Commentary: GPR160 De-Orphanization Reveals Critical Roles in Neuropathic Pain in Rodents (Finally, a Receptor for CART Peptide). Adv Drug Alcohol Res 1, 10012 (2021).

70. G. L. Yosten et al., GPR160 de-orphanization reveals critical roles in neuropathic pain in rodents. J Clin Invest 130, 2587–2592 (2020).

71. L. C. Freitas-Lima et al., GPR160 is not a receptor of anorexigenic cocaine- and amphetamine-regulated transcript peptide. Eur J Pharmacol 949, 175713 (2023).

72. A. Dziedzic, E. Miller, J. Saluk-Bijak, M. Bijak, The GPR17 Receptor-A Promising Goal for Therapy and a Potential Marker of the Neurodegenerative Process in Multiple Sclerosis. Int J Mol Sci 21, (2020).

73. H. J. Yang, A. Vainshtein, G. Maik-Rachline, E. Peles, G protein-coupled receptor 37 is a negative regulator of oligodendrocyte differentiation and myelination. Nat Commun 7, 10884 (2016).

74. D. K. Vassilatis et al., The G protein-coupled receptor repertoires of human and mouse. Proc Natl Acad Sci U S A 100, 4903–4908 (2003).

75. H. Chang et al., Structural basis of oligomerization-modulated activation and autoinhibition of orphan receptor GPR3. Cell Rep 44, 115478 (2025).

76. Y. Liao, G. K. Smyth, W. Shi, The Subread aligner: fast, accurate and scalable read mapping by seed-and-vote. Nucleic Acids Res 41, e108 (2013).

77. Y. Liao, G. K. Smyth, W. Shi, featureCounts: an efficient general purpose program for assigning sequence reads to genomic features. Bioinformatics 30, 923–930 (2014).

78. R. Patro, G. Duggal, M. I. Love, R. A. Irizarry, C. Kingsford, Salmon provides fast and bias-aware quantification of transcript expression. Nat Methods 14, 417–419 (2017).

79. M. I. Love, J. B. Hogenesch, R. A. Irizarry, Modeling of RNA-seq fragment sequence bias reduces systematic errors in transcript abundance estimation. Nat Biotechnol 34, 1287–1291 (2016).

80. M. I. Love, C. Soneson, R. Patro, Swimming downstream: statistical analysis of differential transcript usage following Salmon quantification. F1000Res 7, 952 (2018).

81. J. Feng, T. Liu, B. Qin, Y. Zhang, X. S. Liu, Identifying ChIP-seq enrichment using MACS. Nat Protoc 7, 1728–1740 (2012).

82. Y. Zhang et al., Model-based analysis of ChIP-Seq (MACS). Genome Biol 9, R137 (2008).

83. Q. Wang, et al., Exploring Epigenomic Datasets by ChIPseeker. Curr Protoc 2, e585 (2022).

84. T. C. Murakami et al., An open-source photobleacher for fluorescence imaging of large pigment-rich tissues. Proc Natl Acad Sci U S A 122, e2426628122 (2025).

